# Unjamming overcomes kinetic and proliferation arrest in terminally differentiated cells and promotes collective motility of carcinoma

**DOI:** 10.1101/388553

**Authors:** Andrea Palamidessi, Chiara Malinverno, Emanuela Frittoli, Salvatore Corallino, Elisa Barbieri, Sara Sigismund, Pier Paolo Di Fiore, Galina V. Beznoussenko, Emanuele Martini, Massimiliano Garre, Dario Parazzoli, Ines Ferrara, Claudio Tripodo, Fabio Giavazzi, Roberto Cerbino, Giorgio Scita

**Affiliations:** IFOM, the FIRC Institute of Molecular Oncology, Via Adamello 16, 20139, Milan, Italy; University of Milan, Department of Oncology and Hemato-Oncology, Via Festa del Perdono 7, 20122 Milan, Italy.; Program of Molecular Medicine, European Institute of Oncology, Via Ripamonti 435, Milan, 20141, Italy; Department of Health Sciences, Human Pathology Section, University of Palermo School of Medicine Via del Vespro 129, 90127, Palermo, Italy.; University of Milan, Department of Medical Biotechnology and Translational Med., I-20090 Segrate, Italy

## Abstract

During wound repair, branching morphogenesis and carcinoma dissemination, cellular rearrangements are fostered by a solid-to-liquid transition known as unjamming. The biomolecular machinery behind unjamming, its physiological and clinical relevance remain, however, a mystery. Here, we combine biophysical and biochemical analysis to study unjamming in a variety of epithelial 2D and 3D collectives: monolayers, differentiated normal mammary cysts, spheroid models of breast ductal carcinoma in situ (DCIS), and *ex vivo* slices of orthotopically-implanted DCIS. In all cases, elevation of the small GTPase RAB5A sparks unjamming by promoting non-clathrin-dependent internalization of epidermal growth factor receptor that leads to hyper-activation of endosomally-confined ERK1/2 and phosphorylation of the actin nucleator WAVE2. Physically, activation of this pathway causes highly coordinated flocking of the cells, with striking rotational motion in 3D that eventually leads to matrix remodelling and collective invasiveness of otherwise jammed carcinoma. The identified endo-ERK1/2 pathway provides an effective switch for unjamming through flocking to promote epithelial tissues morphogenesis and carcinoma invasion and dissemination.

## Introduction

Collective motility, a widely recognized mode of migration during embryogenesis, wound repair and cancer^1, 2^, refers to the process of many cells migrating as a cohesive group with a high degree of coordination between neighbouring cells. A complex network of biochemical and physical interactions governs cellular and multicellular motility^2-5^. How cellular and supra-cellular biomechanics and biochemical wiring are integrated and impact onto each other remains, however, largely unexplored.

An emerging framework to interpret these interactions in unifying principles is the notion of cell jamming^6-8^. During tissue growth, cells are rather free to move around, as in a fluid. As density rises, the motion of each cell is constrained by the crowding due to its neighbours. At a critical density, motility ceases and collectives rigidify undergoing a liquid (<u>u</u>njammed)-to-solid (jammed) transition^6-8^, herein referred to as UJT. This transition, which depends on a variety of biophysical parameters such as cell shape variance^9^, intercellular adhesion, cortical tension and single cell motility, is thought to ensure proper development of barrier properties in epithelial tissues, but also to act as a tumour suppressive mechanism^6-8, 10^. The reverse jamming-to-<u>u</u>njamming transition (JUT) might, instead, represent a complementary gateway to epithelial cell migration, enabling tissues to escape the caging imposed by the crowded cellular landscape of mature epithelia^6, 10-12^. Indeed, whereas Epithelial-to-Mesenchymal Transition (EMT) has emerged as the overarching mechanism enabling the dissemination of single tumour cells^13, 14^, invasion by epithelial malignancies (carcinomas) frequently involves the collective migration of cohesive cohorts (nests, sheets, or glandular/tubular structures) of cells into adjacent tissues rather than the scattering of individual carcinoma cells^15, 16^. A number of recent findings supports this hypothesis: i) breast carcinoma frequently disseminate by keeping their epithelial identity, i.e. tight cell-cell interactions and organization into cell cohorts or clusters^17, 18^; ii) circulating cancer cells efficiently seed distant metastasis by forming epithelial cell clusters that maintain cohesive cell-cell interactions, and by doing so display increased metastatic seeding potential^19^; iii) histopathological studies suggest that human invasive ductal breast carcinoma (DCIS) can invade collectively as strands or clusters that retain E-Cadherin-based cellular junctions^20, 21^; iv) late-stage HER2-expressing murine mammary cancers have been shown to undergo kinetic arrest and display reduced metastatic potential as a consequence of increased density and cell packing^22, 23^. These findings further imply that mechanisms capable of overcoming the jammed, kinetically silent state of advanced epithelial malignancies might promote cancer dissemination without the need to invoke changes of cell identity or rewiring of transcriptional programs. However, how cells control the JUT is unclear.

Membrane trafficking circuitries have emerged as pivotal in regulating the duration, intensity, and spatial distribution of signals, thereby contributing to pathway specificity^24, 25^, with a primary role on cell migration plasticity and on the mechanics of cell-cell interactions^26-28^. Consistent with the above notion, we recently found that endocytic circuitries controlled by RAB5A, a master regulator of early endosomes necessary to promote a proteolytic, mesenchymal program of individual cancer cell invasion^29, 30^, have a dramatic impact on the mechanics and dynamics of multicellular, normal and tumorigenic, cell assemblies^31^. Elevation of RAB5A levels is sufficient to re-awaken the motility of otherwise jammed and kinetically arrested epithelial monolayers^31^. RAB5A does so by increasing monolayers stiffness, cell-cell surface contact and junctional tension, while concomitantly accelerating the turnover of junctional E-cadherin^31^. RAB5A further promotes millimetres-scale, ballistic locomotion of multicellular streams by augmenting the extension of oriented and persistent RAC1-driven, protrusions^31^. These effects combine to endow monolayers with a flocking fluid mode of motion, which is explained in numerical simulations in terms of large scale coordinated migration and local unjamming, driven by an increased capability of each individual cell to align its velocity to the one of the surrounding group^31-33^. Molecularly, impairing endocytosis, macropinocytosis or increasing fluid efflux abrogated RAB5A-induced collective motility, suggesting that perturbations of trafficking processes impacting on different signalling and biomechanical pathways are necessary for the JUT. However, the molecular nature of these endocytic-sensitive pathways remains to be identified. Even less clear is whether JUT occurs in relevant physiological setting and whether is hijacked by dense and jammed carcinoma to promote their collective dissemination.

Here, using kinematic and biochemical analysis of jammed monolayers dynamics, we showed that enhanced epidermal growth factor receptor (EGFR) internalization through non-clathrin dependent routes leads to endosomal ERK1/2 hyper-activation and phosphorylation of the branched actin nucleator, WAVE2^34, 35^. This endo-ERK1/2 pathway, in turn, is critical to promote a transition to a flocking-liquid mode of collective motility. Importantly, this pathway is also sufficient to overcome kinetic and proliferation arrest of fully differentiated normal mammary cysts in 3D, and to initiate bud morphogenesis. In DCIS models, instead, endocytic-mediated unjamming endows tumour spheroids embedded into thick collagen matrix with a striking and coordinated circular angular motion (CAM), that promotes matrix remodelling and collective local invasion, recapitulating what is observed in DCIS foci orthotopically implanted into recipient mice. We propose that the EGF-dependent activation of endosomal ERK1/2 as the first identified molecular route to the JUT via flocking, sufficient to overcome the kinetic and proliferation arrest of terminal differentiated epithelial cells, and to promote collective invasive programs of jammed breast carcinoma.

## Results

### Endocytic reawakening of motility depends on EGFR activation and is caused by alterations of EGFR trafficking

RAB5A is deregulated in breast cancer (BC)^30, 36^. By focusing on early stages of BC progression, we found that RAB5A expression was variably low in malignant cells of densely-packed and jammed ductal carcinoma in situ (DCIS) foci and increased at foci of DCIS with invasion or in overt infiltrating cancers (Fig.1A). Additionally, its elevated expression is detected in various malignant, aggressive BC cell lines (Fig. 1B), and, more relevantly, correlates with worse relapse free probability in various BC subtypes (Fig. S1). We employ doxycycline-inducible RAB5A-MCF10A-line to induce the expression of the GTPase to levels that mimic those encountered in human DCIS (Fig. 1B), and study the molecular mechanisms of RAB5A impact on multicellular kinematics and jamming transition^31^. Fully confluent, dense epithelial monolayers of normal mammary MCF10A cells are locked into a jammed, solid state characterized by a full kinetic arrest (Ref.^31^ and Movie S1and S4). Doxycycline-mediated induction of RAB5A is sufficient to reawaken the motility by promoting large cellular streams (Fig. 1C-D and Movie S1-4 and ref.^31^). Particle Image Velocimetry (PIV) analysis was used to capture the kinematic of cell locomotion in jammed epithelia. As previously reported^31^, RAB5A expression enhanced robustly the root mean square velocity (*v*_*RMS*_) of the cells, and promoted millimetres-scale cell coordination as revealed by calculating the correlation length, *L* _*corr*_, as the width of the correlation function 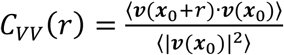 of the (vectorial) velocity *v*(*x*_0_), whose typical width provides an estimate of the velocity correlation length^37^. We also quantified cellular motion using the Mean Square Displacement (MSD) over a given time interval, δt, averaged over many cells in several optical fields (see Methods), which was used to extract an estimate of the persistence length, L_*pers*_^37^. The latter corresponds to the distance travelled by a cell at constant velocity before the direction of its motion becomes uncorrelated with the initial one. RAB5A-expression promoted persistent and ballistic collective motion over a distance larger than 700 μm, consistent with monolayers acquiring a flowing, liquid mode of motion. Removal of EGF, required for proliferation and single cell motility of MCF10A^38^, or addition of AG1478, an inhibitor of EGFR kinase^39^, arrested the flowing mode of motion induced by RAB5A (Fig. 1C-D and Movie S1 and S2 and S4), reduced *v*_*RMS*_, *L*_*corr*_, and *L*_*pers*_ to values seen in control cells. These treatments further impacted on the uniformity of the migration pattern captured by local alignment (*a*) of the velocity vector with respect to the mean velocity, which varies between +1 and -1 when it is parallel or antiparallel to the mean direction of migration, respectively (Fig. 1D). We further corroborated these results using EGFP-H2B control and RAB5A-expressing cells to visualize directly nuclear cell displacement within the epithelial collective (Movie S3). Finally, similar EGF–dependency of collective motion was also observed in jammed keratinocyte monolayers, HaCat, (Ref.^31^ and not shown) and in oncogenically-transformed MCF10A variants, MCF10.DCIS.com (see below Fig. S5A and Movie S16).

**Figure 1.**
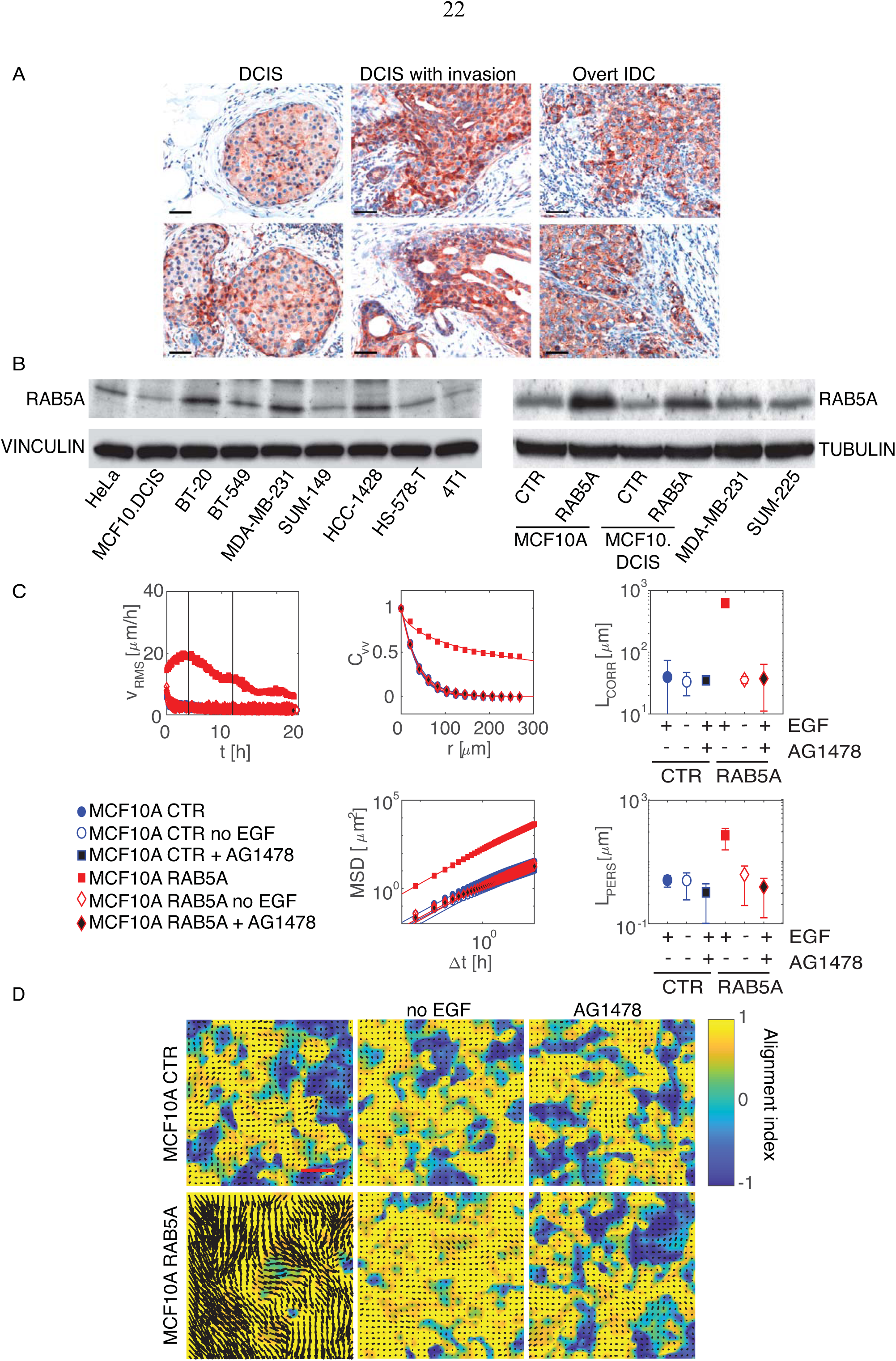
Endocytic reawakening of motility is strictly dependent on EGFR activation. **A**.Representative immunohistochemical staining of RAB5A on human ductal breast carcinoma in situ (DCIS) foci, DCIS with invasive components, and overtly infiltrative breast cancer foci showing the heterogeneous expression of RAB5A in DCIS foci and its increase in invasive areas. Scale bar, 100 μm. Please note the heterogenous and elevated expression of RAB5A in human IDC vs DCIS. **B**.Doxycycline-treatment of MCF10A and MCF.DCIS.com engineered to express RAB5A in an inducible fashion increases the level of the protein, mimicking those found a variety of BC lines shown in the lower panels. Immunoblotting of total cell lysates with the indicated abs. Tubulin and Vinculin were used as loading control. **C**.PIV analysis of motion of doxycycline-treated control and RAB5A-MCF10A cells seeded at a jamming density and monitored by time-lapse microscopy in the presence or the absence of EGF (Movie S1 and 3) or after treatment with the EGFR inhibitor AG1478 (Movie S2). Vertical lines indicate the time interval used for the analysis of the following motility parameters: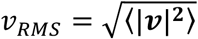 root mean square velocity (representative of > 5 independent experiments); *C*_*vv*_; velocity correlation functions as function of the distance *x,j*. The correlation function is evaluated in the time window comprised between 4 and 12 h during which a peak in *v*_*RMS*_ is observed. In all cases, *C*_*vv*_ is well fitted to a stretched exponential decay, with stretching exponent γ decreasing from 0.91±0.06 (control) to 0.62±0.04 (RAB5A). *L*_*corr*_: correlation lengths whose width provide an estimate of the size of group of cells moving in coordinated fashion, which in RAB5A-expressing cells is close to 0.78±0.3 mm, (corresponding to more than 50 cell diameter), whereas in control or after EGF-deprivation or treatment with AG1478 is around 44±6 μm (1-to-2 cell diameter). *MSD*: mean square displacements obtained by numerical integration of the velocity maps over a given time interval, δt. In all cases, for short times *MSD* displays a quadratic scaling *MSD* ≅ (*u*_0_Δ*t*)^2^, which is indicative of a directed ballistic motion, although with dramatically different characteristic velocities (*u*_0_=36 μm/h for RAB5A, *u*_0_< 7 μm/h for the control or w/ o EGF or in the presence of AG1578). At later times, a transition to a diffusive-like regime characterized by a scaling exponent close to 1 is observed. By fitting the MSD curves with a model function (continuous lines-see methods), we extracted an estimate of a persistence lengths, *L*_*pers*_, which in RAB5A is around 450±50, while in all other conditions is less than 65±3.1. Data were obtained by analysis of at least 5 movies/experimental conditions out of at least 4 independent experiments. **D**.Snapshots of the velocity field obtained from PIV analysis of doxycycline-treated control (Ctrl) and RAB5A-MCF-10A cells seeded at jamming density in the presence or the absence of EGF or treated with the EGFR inhibitor, AG1478, and monitored by time-lapse microscopy (Movie S4). The colour-map reflects the alignment with respect to the mean instantaneous velocity, quantified by the parameter *a*(*x*) = (*v*(*x*). *v*_0_)/(|*v*(*x*)||*v*_0_|). *a* = 1(−1), the local velocity is parallel (antiparallel) to the mean direction of migration (not shown). Scale Bar, 100 μm.

This finding is consistent with the possibility that alterations of endosomal biogenesis caused by RAB5A^40^ and leading to reawakening of collective motion^31^ might specifically perturb EGFR cellular distribution, trafficking or signalling. We set out to test these possibilities. Firstly, we showed that the total protein, but not mRNA levels of EGFR were significantly reduced following induction of RAB5A expression (Fig. 2A). The fraction of phosphorylated EGFR was, instead, unexpectedly increased (Fig. 2A, Table), suggesting an impact of RAB5A on cellular distribution and trafficking of this receptor. Consistently, immunofluorescent analysis revealed that RAB5A-expressing cells display a marked reduction of cell surface EGFR, as detected in non-permeabilized cells, accompanied by a sizable increase of intracellular EGFR, which accumulates in EEA1 positive vesicle (Fig. 2B-C). Measurements of the absolute number of surface EGFR using ^125^I-EGF binding corroborated the immunofluorescence (IF) data (Fig. 2D). MCF10A are grown in the presence of saturating dose of EGF (20 ng/ml), which binds and activates EGFR, promote its rapid internalization and subsequent lysosomal degradation. Thus, RAB5A may perturb EGFR cellular distribution by enhancing its internalization, trafficking and degradation. If this were the case removal of EGF or inhibition of its kinase activity should restore EGFR surface and intracellular distribution to levels seen in control cells. We verified this prediction by IF analysis (Fig. 2E-G), and by determining the number of EGFR molecules on cell surface (Fig. 2H).

**Figure 2.**
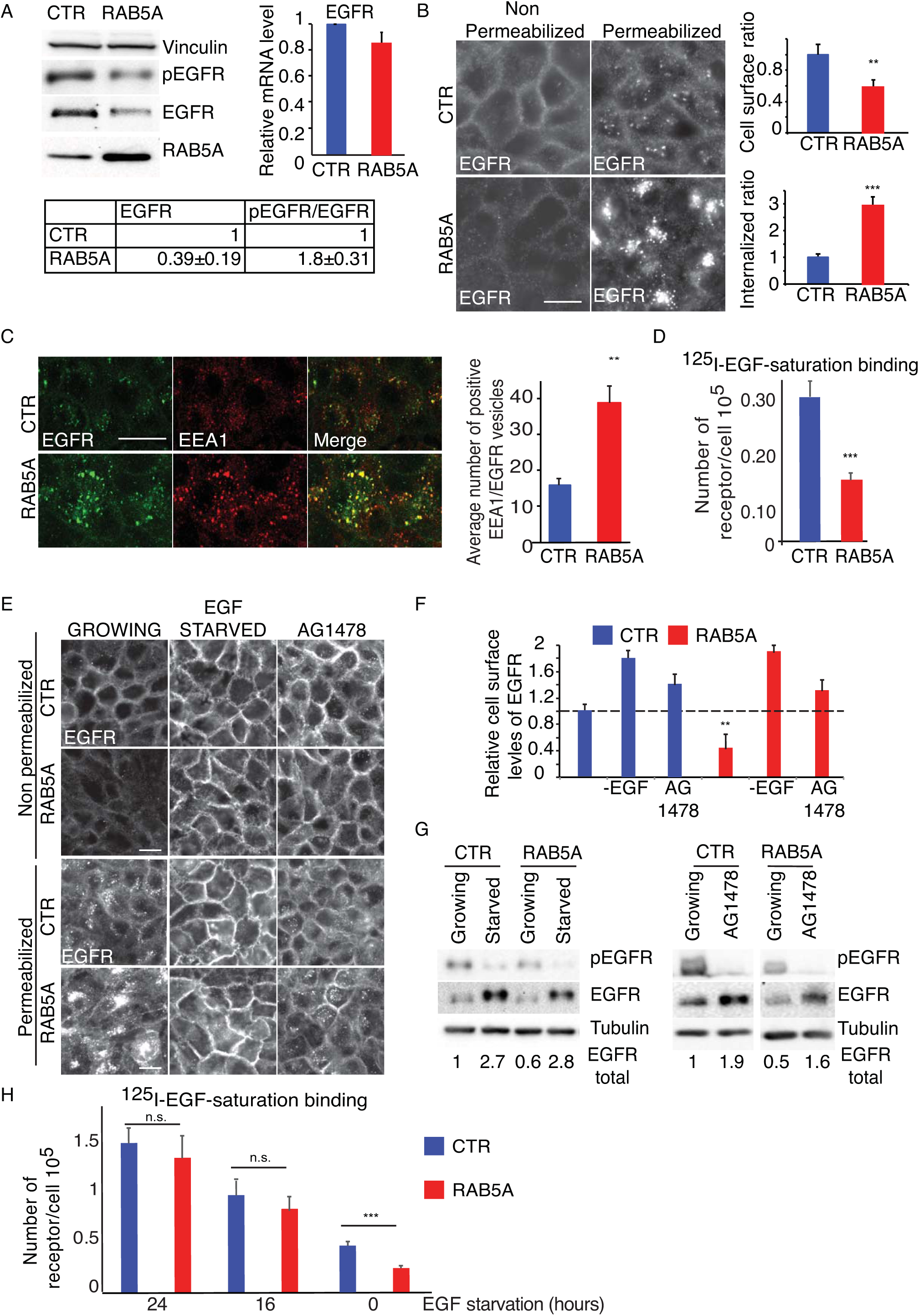
RAB5A alters EGFR cellular distribution, trafficking and stability. **A**.Total cellular proteins (left) and EGFR mRNA levels (right) of control (CTR) and RAB5A-MCF10A seeded at jamming density and detected by immunoblotting or qRT-PCR, respectively. At the bottom, quantification of total EGFR and of the ratio of phosphorylated/total EGFR from the immunoblotting is shown. Data are the relative ratio (mean±SD, n=5 independent experiments) with respect to control, obtained after normalizing each intensity values to Vinculin. The levels of EGFR mRNA are expressed relative to control (mean±SD, n=5 independent experiments) after normalizing to GAPDH. **B**.Control and RAB5A-MCF10A cells seeded at a jamming density were fixed and either permeabilized or non-permeabilized with 0.1% Triton X100 before staining with anti-EGFR ab. Data are the relative ratio (mean ± SD) with respect to control of total cell surface or internalized EGFR signals (n = 100 cells out of at least 3 independent experiments) normalized to cell number. Scale Bar, 20 μm.**p< 0.01, ***p<0.005. *P* values were calculated using each-pair Student’s t-test. **C**.Representative images of control and RAB5A-MCF10A cells seeded at a jamming density, fixed and stained with the indicated abs. Data are the mean±SD of positive EEA1and EGFR vesicles/cells (n>150 out of 3 independent experiments). ** p< 0.01, Student’s t-test. Scale Bar, 20 μm **D**.Number of EGFR/cell of control and RAB5-MCF10A seeded at jamming density measured by^125^I-EGF saturation binding after subtracting unspecific background counts (see methods for details). Data are the mean±SD of triplicate measurements of a representative experiment. *** p<0.005. *P* values were calculated using each-pair Student’s *t* test. **E-G**. Representative images (**E**) of control and RAB5A-MCF10A cells seeded at a jamming density, which were either deprived of EGF for 24 h or treated with AG1478 before fixation and staining as described in B. Scale Bar, 20 μm. Data (**F**) are the relative ratio (mean ± SD) with respect to control of total cell surface EGFR (n>120 cells in 2 independent experiments) normalized to cell number. ** p<0.05, each-pair Student’s t*-*test versus control. Immunoblots (**G**) showing the efficacy of EGF deprivation and EGFR inhibition on total and phosphorylated EGFR levels using the indicated abs. Each band of total EGFR was quantified and its mean intensity value is shown. The experiment is representative of at least 5 independent ones with similar outcome. **H.** Number of EGFR/cell of control and RAB5-MCF10A seeded at jamming density measured by ^125^I-EGF saturation binding after various time of EGF starvation as described in D. Data are the mean±SD of triplicate measurements of representatives experiments. * p<0.05. *P* values were calculated using each-pair Student’s t-test.

Intracellular accumulation of EGFR might be the results of elevated internalization or reduced recycling. In the former case, although clathrin-mediated endocytosis (CME) represents the best-characterized internalization route of EGFR into cells^41^, it can also occur through non-clathrin endocytosis (NCE), depending on growth conditions and cellular context^42-45^. At a low epidermal growth factor (EGF) dose (1 ng/ml), EGFRs are primarily internalized by CME and recycled back to the plasma membrane (PM)^44^. For large physiological EGF concentrations (20 to 100 ng/ml), NCE is activated in parallel to CME. EGFRs entering via NCE (~40%) are predominantly trafficked to the lysosome for degradation^44, 45^. To test whether RAB5A expression influences any of these entry routes, we measured the rate of internalization of ^125^I-EGF at low and high concentrations. RAB5A expression significantly increased the endocytic rate constant (Ke) at high (30 ng/ml), but not at low (1 ng/ml) EGF concentrations, suggesting a specific impact on NCE (Fig. 3A). Using a similar approach, we also measured recycling rates of EGFR, which were not significantly altered by elevation of RAB5A (Fig. 3B). Following doxycycline induction of RAB5A expression, we further monitored the total levels of EGFR, which were slowly, but progressively decreased over time consistently with the augmented NCE internalization into endocytic compartments (Fig. 3C). Collectively, these findings indicate that RAB5A promotes EGFR NCE internalization routes, likely leading to increased endosomal EGFR and, possibly, to the re-awakening of collective motion.

**Figure 3.**
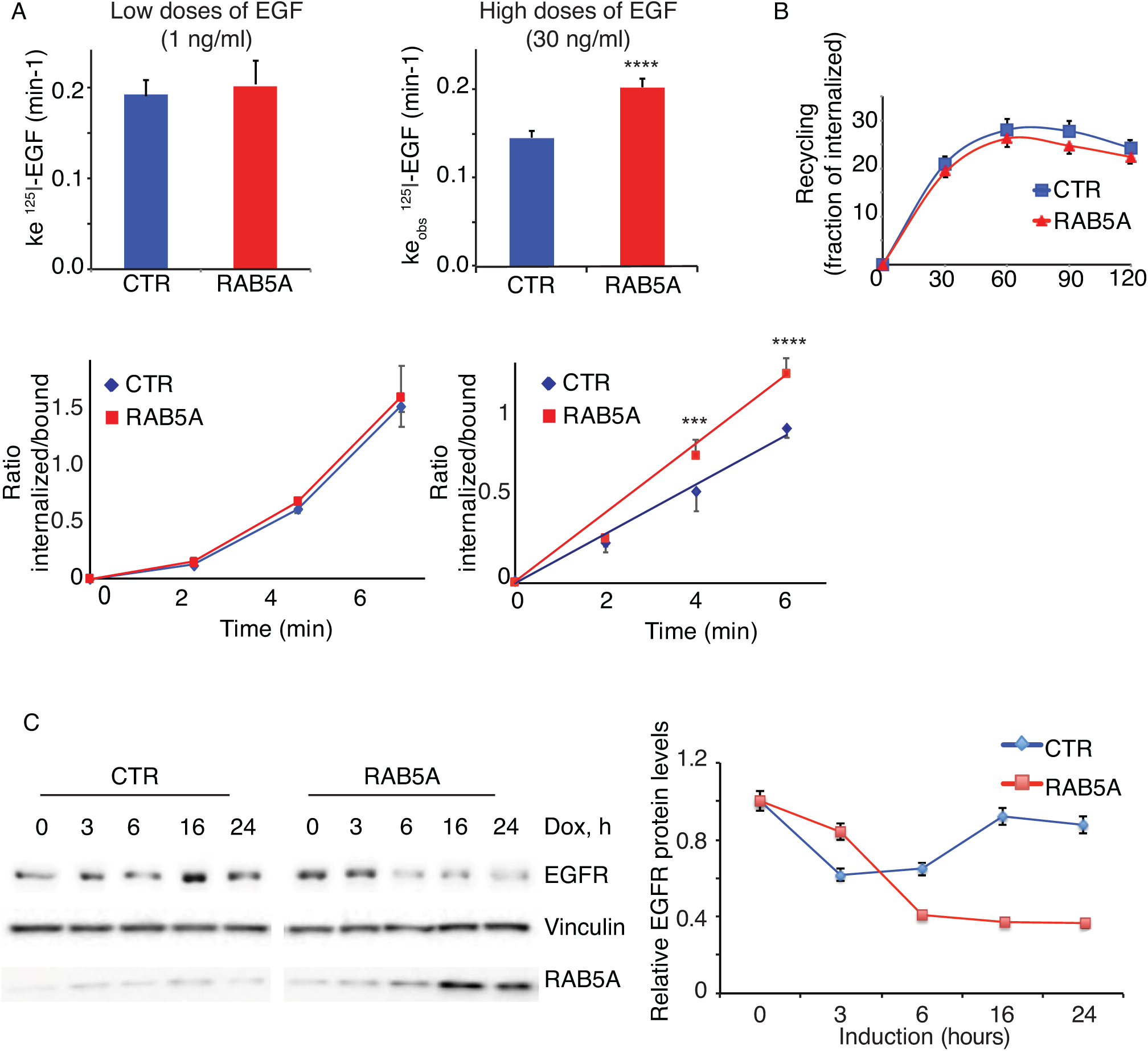
RAB5A enhances internalization of EGFR at high doses of ligand. **A**.EGFR internalization kinetics in control and RAB5A-MCF10A cells seeded at jamming density was measured using ^125^I-EGF at low (1 ng/ml) or high (30 ng/ml) concentrations. Results are expressed as the effective or apparent internalization rate constants at low and high EGF doses, respectively (Ke, upper panel). A representative kinetic of the ratio of ^125^I-EGF internalized/bound is shown at different time points and is expressed as the mean±SD (n=3 out of 5 independent experiments). ****p <0.0005. *P* values were calculated using each-pair Student’s t*-*test. **B**.Control or RAB5A-MCF10A cells seeded at jamming density were incubated with ^125^I-EGF (30 ng/ml) for 15 min at 37 °C. Recycling of ^125^I-EGF at the indicated time points was estimated as described in Methods. Data are the mean ± SD (n = 3 replicates in a representative experiment). **C.A** representative kinetic of total EGFR levels as function of time after treatment with doxycycline in control or RAB5A-MCF10A cells. At the indicated time, total cellular lysates were immunoblotted with the indicated abs. The EGFR levels relative to control and normalized to Vinculin, used as loaded control are shown. Data are the mean±SD (n=5 independent experiments). Representative blots are shown.

### Activation of endosomal ERK1/2 is a molecular route to unjamming via flocking

EGFR signalling and trafficking are strictly interdependent^46^. For example, the detection of phosphorylated receptors and signalling adaptors in endosomes indicated that signalling is initiated at the plasma membrane but continues in endosomes^47-49^. Albeit recent work challenged this concept^50, 51^, quantitative high-resolution FRET microscopy demonstrated that phosphorylated EGFR can be packaged at constant mean amounts in endosomes, which were proposed to act as signalling quanta-like platforms^52^. As a consequence, altering the size and number of endosomes directly affected the amplitude and duration of EGFR signalling.

Hence, we tested whether any of the canonical EGFR downstream pathways is altered following RAB5A expression. We found that while phosphorylated AKT and p38 levels were not significantly altered in confluent cells, phosphorylated ERK1/2 was elevated and long lived in RAB5A (Fig 4 A- B), but not in RAB5B or C expressing cells (Fig. S2A-B). Notably, RAB5B and C were very inefficient in reawakening collective motion in jammed monolayers (Fig. S2C and Movie S5). We corroborated this finding by testing in situ, through IF, the levels of phosphorylated ERK1/2 in intact monolayers formed by mixing control and EGFP-H2B-RAB5A-expressing cells (Fig. 4C). ERK1/2 has also been reported to be activated in a temporally distinct “two waves” fashion after wounding that propagate in epithelial sheet controlling collective motion^53^. We found that RAB5A-MCF10A cells, which display accelerated wound migration speed, display a robust increase in ERK1/2 wave amplitude (Fig. S2D). Pharmacological inhibition of the ERK1/2 using PD0325901 that targets the upstream MEK kinase^54^, abrogated flocking mode of locomotion of RAB5A-monolayers by reducing *v*_*RMS*_, *L*_*corr*_ and *L*_*pers*_ to control levels (Fig. 4D and Movie S6). RAB5A-mediated elevation of ERK1/2 (Fig. 4E) was inhibited by treatment of MCF10A with AG1478 or Dynasore, a small molecule impairing dynamin pinchase activity^55^. These treatments also impeded reawakening of collective motion in jammed epithelia (Fig. 1, Movie S2, Movie S7 and ref^31^). Similar results were also obtained by silencing the expression of Dynamin 2, the only dynamin isoform expressed in MCF10A (Fig. S3A and Movie S8). The sum of these findings indicates that RAB5A elevation specifically enhances endosomally-compartmentalized ERK1/2 signalling. To directly test this possibility, we generated FRET EKAREV-ERK1/2 sensor^56^, which was targeted to endosomes by appending to its C-terminus the FYVE domain of SARA protein^57^ (Fig. 4F). The FYVE-ERK1/2-EKAREV-FRET sensor was, indeed, found on EEA1-positive endosomes (Fig. 4G), which were increased in size and dimension following RAB5A expression^31^. Removal of EGF or treatment of cells with PD0325901, significantly impaired FRET efficiency validating the biological relevance of the sensor (Fig. 4H). More importantly, RAB5A-expressing cells displayed elevated endosomal ERK1/2 FRET efficiency as compared to control monolayers (Fig. 4H). We further showed that global (or plasma membrane associated) elevation of ERK1/2 phosphorylation brought about by the expression of a constitutively activated MEK-DD stably expressed in MCF10A was insufficient to reawaken motility in jammed monolayers (Fig. S3B-C and Movie S9), reinforcing the notion that ERK1/2 must be activated in endosome to promote unjamming. Notably, inhibition of the late endosomal ERK1/2 scaffold, MP1or p14^58, 59^, had no impact on RAB5A mediated ERK1/2 hyper-activation, nor on collective motion (Fig. S3D-F and Movie S10) suggesting that other, yet to be identified molecular determinants/scaffolds mediate early endosome compartmentalization of ERK1/2 signalling.

**Figure 4.**
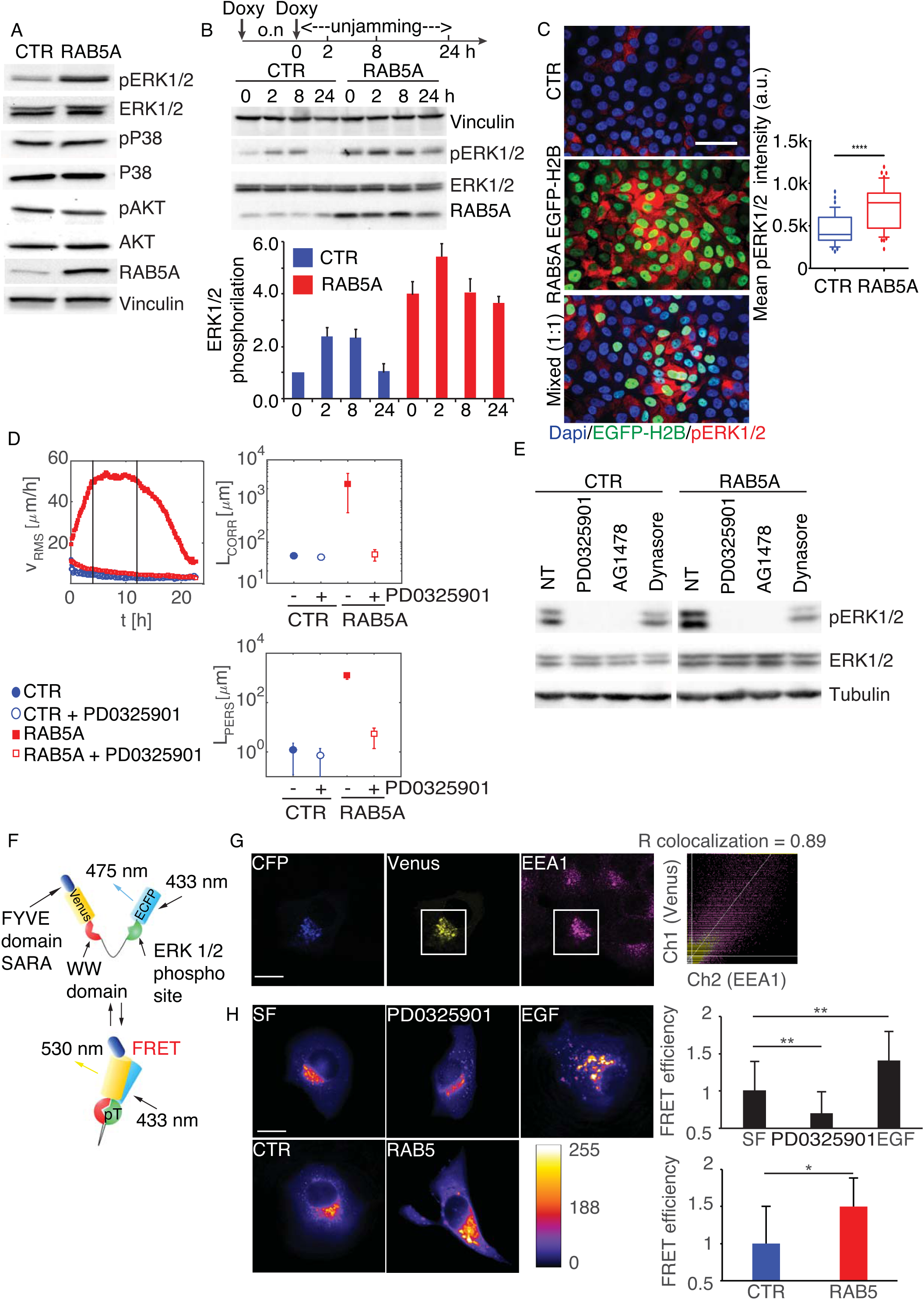
RAB5A-induced, EGFR-dependent endosomal ERK1/2 activity is required for flocking locomotion in epithelial monolayer. **A**.Total cellular lysates of control and RAB5A-MCF10A cells seeded at jamming density and treated with doxycycline for 16 h to induced RAB5A expression were immunoblotted with the indicated abs. **B**.Total cellular lysates of control and RAB5A-MCF10A cells seeded at jamming density were treated with doxycycline overnight (o.n.). The morning after, the media was replenished and cells were either monitored by time lapse to follow their kinematics or lysed at various time point in order to follow the phosphorylation status by immunoblotting of ERK1/2, coincidentally with the expression of RAB5A and the reawakening of locomotion (not shown), as indicated in the experimental scheme reported above. The ratio of phosphoERK1/2/totalERK1/2 is plotted below and expressed as mean±SD (n=3 independent experiments). **C**.Doxycycline-induced control and EGFP-H2B-RAB5A cells were seeded at jamming either alone or mixed at a 1:1 ratio (Mixed) fixed and stained against phosphorylated ERK1/2 or processed for epifluorescence. The mean±SD of the intensity of pERK1/2 is shown in the box plots (n=200 cells in at 3 independent experiments). Box and whisker: 10-90 percentiles. Outliers are plotted as bubbles and medias are horizontal lines in the boxes. ****p<0.0001. P value was calculated Student’s t*-*test. Scale Bar, 50 μm. **D**.Doxycycline-treated control and RAB5A-MCF10A cells seeded at jamming density were incubated with vehicle or PD0325901 (1 μM), a MEK inhibitor, 1 h before starting time-lapse recording (Movie S6). PIV analysis was applied to extract: root mean square velocity *v*_*RMS*_, plotted as a function of time, correlation length *L*_*corr*_ and persistence length *L*_*pers*_. Data are the mean±SD (n=5 movies/ conditions out of 3 independent experiments). **E**.Total cellular lysate of doxycycline-treated control and RAB5A-MCF10A cells seeded at jamming density treated with PD0325901, or AG1478, or Dynasore (80 μg/ml) or vehicle as control were immunoblotted with the indicated abs. Data are representative of 4 experiments with similar outcome. **F**.Scheme of the Endo-ERK1/2-FRET sensor. The FYVE domain of SARA protein is appended at the C-terminus of an EKAREV construct^56^ carrying from C-terminus to the N-terminus: Venus, the domain binding WW phosphopeptide, a 72-Gly linker, a ERK1/2 Serine-containing substrate (sensor domain) and ECFP. **G**.Endo-ERK1/2-FRET sensor localized into EEA1-positive early endosomes. Representative images of Endo-ERK1/2-FRET transfected MCF10A cells stained with anti-EEA1. The extent of colocalization between the Endo-ERK1/2-FRET sensor and EEA1 is shown on the right as Pearson Correlation coefficient. Scale Bar, 20 μm ***H***. *Upper panels*, Endo-ERK1/2-FRET transfected control MCF10A cells were either serum starved (SF) for 24 h or treated with PD0325901 or incubated with EGF (20ng/ml), then fixed and processed for detection of FRET efficiency as described in methods. FRET efficiency normalized to control, serum free MCF10A cells are the mean±SD (n=55 cells/experimental condition in three independent experiments). *Bottom panels*, Endo-ERK1/2-FRET transfected control or RAB5A-MCF10A cells were processed for epifluorescence as described above. FRET efficiency normalized to the value of control cells is expressed as mean±SD (n=75 cells/experimental condition in 3 independent experiments). Scale Bar, 20 μm *p<0.05, ** p<0.01. P value were calculated using Student’s t-test.

What are the molecular substrates that endosomal ERK1/2 activates to promote collective motion? RAB5A-expressing, unjammed monolayers move in a highly ballistic, directed fashion by extending cryptic and oriented lamellipodia^60^ underneath neighbouring cells^31^. The latter structures are dependent on RAC1, which activates branched actin polymerization of the pentameric WAVE2 complex^61^. The key component of this complex, WAVE2, a nucleation promoting factor, must be phosphorylated by ERK1/2 on multiple serine residues, among which S351 and S343, to be activated and to control protrusion initiation and speed^62, 63^. Consistent with the latter finding, using phosphospecific antibody, we found that RAB5A expression increased the phosphorylation of S351 and of S343 of WAVE2 (Fig. 5A) in an ERK1/2, EGFR and Dynamin-dependent manner (Fig. 5B). Additionally, by monitoring the dynamics of cells mosaically-expressing EGFP-LifeAct, we found that pharmacological inhibition of ERK1/2 impaired the formation of cryptic lamellipodia (Fig. 5C and Movie S11). Thus, RAB5A promotes endosomal, EGFR-dependent ERK1/2 signalling leading to hyper phosphorylation and activation of WAVE2, and the formation of persistent lamellipodia that contribute to reawakening of collective motion.

**Figure 5.**
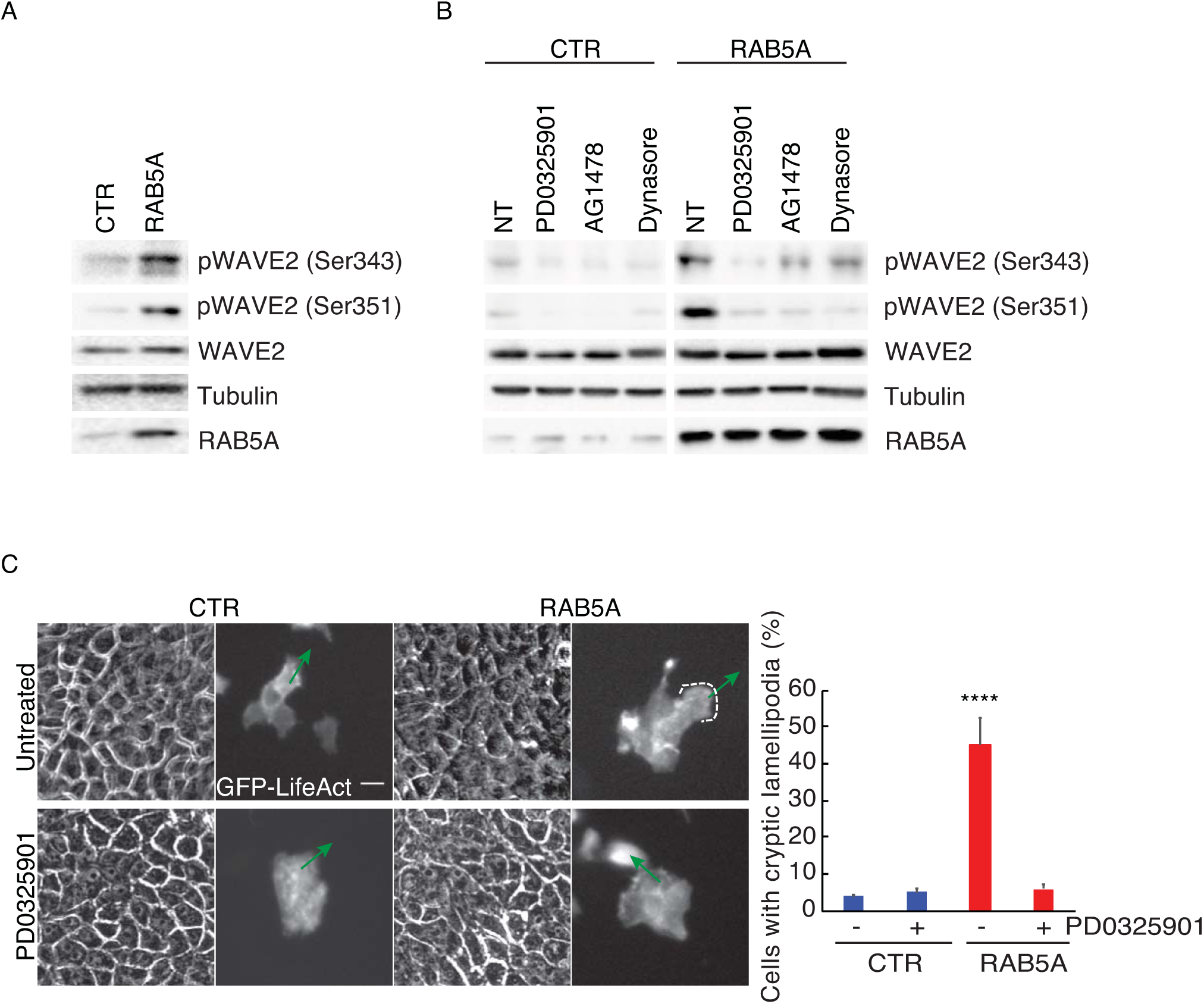
Endosomal ERK1/2 activation leads to phosphorylation of WAVE2, which is required for cryptic lamellipodia extension. **A-B**. Total cellular lysate of doxycycline-treated control and RAB5A-MCF10A cells seeded at jamming density (A) or treated (B) with either vehicle (NT) or with PD0325901, or AG1478, or Dynasore were immunoblotted with the indicated abs. **C.** Still phase-contrast and fluorescent images of cryptic lamellipodia in control and RAB5A-MCF-10A monolayers composed of mosaically GFP-LifeAct-expressing (green):non-expressing cells (1:10 ratio) monitored by time-lapse microscopy (Movie S11). Green arrows indicate the orientations of protrusions. Scale bars, 20 μm. Right plot: proportion of cells with lamellipodium. Data are the mean**±**SD. (n=65 cells/conditions from 4 independent experiments) ****p<0.0001, Student’s t-test.

### RAB5A-mediated unjamming overcomes kinetic and proliferation arrest in terminally-differentiated mammary acini

To explore the biological consequence of RAB5A–induced endo-ERK1/2 axis in more relevant physiological 3D processes, we exploited the well-established ability of MCF10A cells to recapitulate mammary gland morphogenesis when grown in 3D on top of Matrigel plugs in Matrigel-containing media^38^. Under these conditions, cells generate filled spheroid that within 14-to-21 days undergo a full differentiation program, giving raise to apico-basally polarized (Fig. S4A), kinetically and proliferation-arrested hollow cysts^38^. We employed mCherry-H2B-expressing control and RAB5A-cells to monitor kinematics of differentiated cysts treated with doxycycline to induce transgene expression (Fig. 6A). Cells in control differentiated acini display a limited motility and were, as expected, locked in jammed, kinetically-arrested states. The expression of RAB5A reawakened motility by triggering a striking circular angular rotational mode of motion with cells within the cysts migrating in an apparent highly-coordinated fashion (Fig. 6B and Movie S12). We applied a custom PIV analysis to evaluate the tangential velocity field associated with the cellular motion, from which we extracted the relevant kinematic parameters, like the root mean square velocity *v*_*RMS*_ and the rotational order parameter *ψ* (see Methods). The latter, which can vary in the range [0, 1], captures the uniformity of collective motion (see also Methods): *ψ* = 1 corresponds to a rigidly rotating sphere while, in the absence of coordinated motion, one expects *ψ* ≅ 0. Control acini display barely detectable *v*_*RMS*_, while the order parameter Ψ was constantly below 0.2 (Fig. 6C and Movie S12-13). In RAB5A-expressing cysts, we observed a marked elevation of *ψ*, which reached value close to 1 in correspondence of the largest values of *v*_*RMS*_, reflecting the acquisition of collective angular motion (CAM) (Fig. 6C and Movies S12-S13). We exploited this finding to assess whether RAB5A-induced reawakening of motility in 3D cysts was dependent on the same key determinants controlling 2D locomotion using various inhibitors. Indeed, impairing EGFR activity, ERK1/2 phosphorylation and dynamin-endocytosis effectively reduced *v*_*RMS*_ and the order parameter to control levels (Fig. 6C-table and Movies S14). We further tested whether RAB5A promoted similar microscopic changes as the ones seen in 2D unjammed monolayers^31^. Specifically, we analysed the morphology and distribution of E-cadherin junction both by IF and phosphorylated ERK1/2. We observed that cells in RAB5A-cysts display straight and compact junctions and elevated phosphoERK1/2 (Fig. S4B). The junctional features likely account for the large scale, coordinated motility of RAB5A acini. We also noticed that the induction of the expression of RAB5A in the initial phase of cystogenesis reduced the number of acini, but the ones remaining were significantly larger in size (Fig. 6D and Movie S15), and did not undergo proliferation arrest, like control cysts do. Indeed, whereas in control acini we detected no Ki67-positive, proliferating cells after 14 days in overlaid cultures, a sizeable fraction of RAB5A cysts kept on proliferating under conditions in which the number of apoptotic cells was, instead, comparable to that of control acini (Fig. S4D). We investigated this phenotype further by adding doxycycline to induce the expression of RAB5A at the end of the morphogenetic process, when acini have ceased proliferation and motility to complete the differentiation program^38^. RAB5A expression reawakened not only cell motility (see above) but was also sufficient to overcome proliferation arrest (Fig. 6F), in a strict ERK1/2-dependent manner (Fig. 6F).

**Figure 6.**
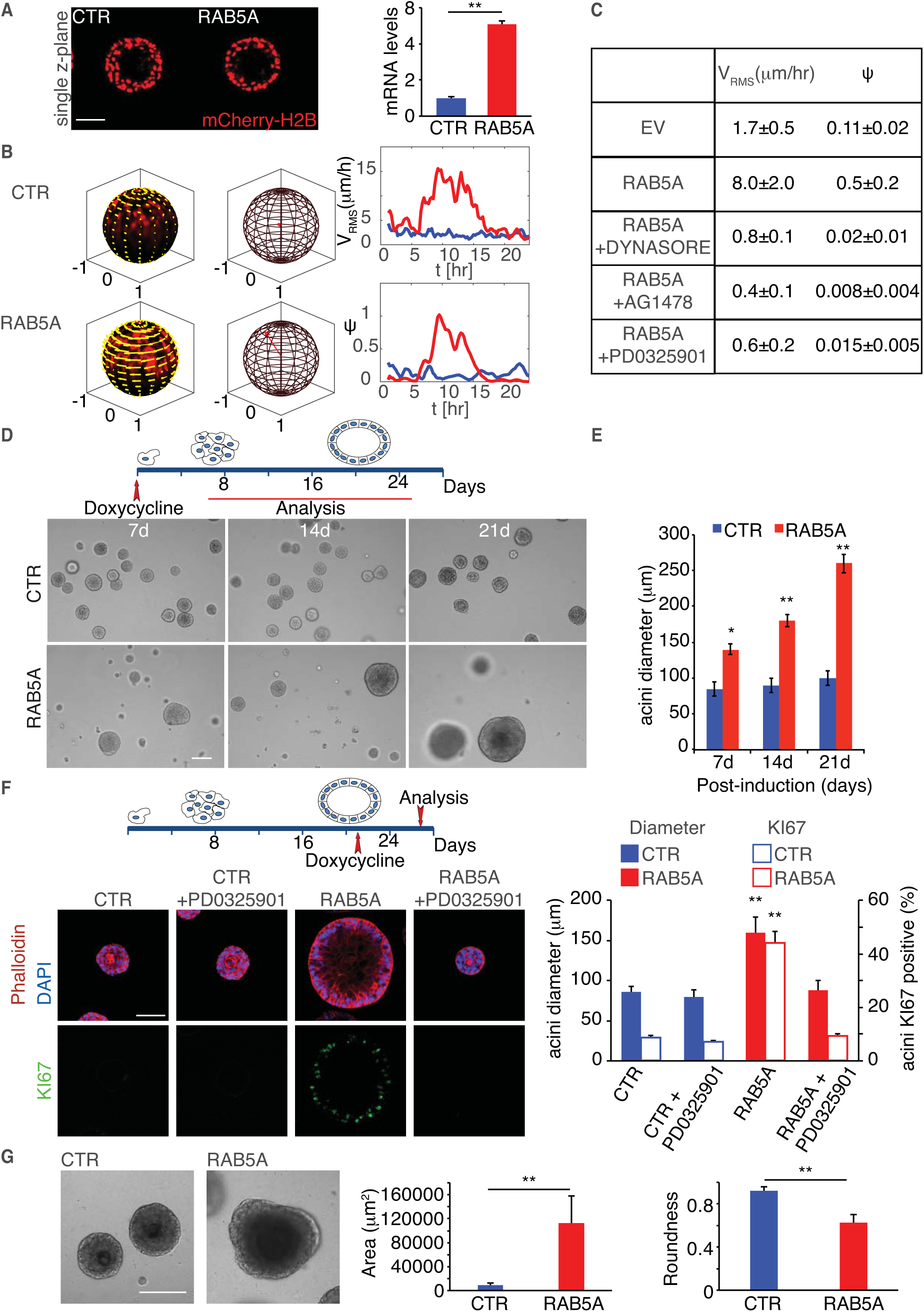
RAB5A-mediated unjamming overcomes kinetic and proliferation arrest in terminal differentiated mammary acini. **A-B**. Control and RAB5A-MCF10A-expressing mCherry-H2B cells were grown overlaid on top of Matrigel plugs. Between 14 and 21 days, cells formed fully differentiated hollow acini. At this stage, Doxycycline was added and the kinematic of the 3D acini was monitored by confocal times lapse for 24 h (Movies S12). Representative images of single Z planes are shown. RAB5A induction was verified by QRT-PCR, expressed relative to control after normalizing to GAPDH. The data are the relative level of gene expression compared to control expressed as mean ± SD (n=3 independent experiments). Scale Bar, 50 μm. In **B**, snapshots of the tangential velocity field at *t* = 10 h (indicated by yellow arrows) obtained from PIV analysis are shown, overlaid on radial projection of the acini onto a unit spherical surface (see also Movies S13). The direction of the red arrow shown in the middle panel is parallel to the instantaneous total angular momentum *l* and provides the orientation of the instantaneous axis of rotation, while its length is equal to the instantaneous order parameter Ψ. On the right: time evolution of the root mean square velocity *v*_*RMS*_ and of the rotational order parameter Ψ (see text and Methods for details). The data are representative of 4 movies in 3 independent experiments. **C**. Doxycycline-treated control and RAB5A-MCF-10A mcherry-H2B-expressing MCF10A acini were treated with the indicated vehicle or the indicated inhibitor and monitored by confocal time-lapse microscopy for 24 hr (Movie S14). Average values of *v*_*RMS*_ and of Ψ, calculated over the time window comprised between 4 and 12 h, are reported. Values are from 5 movies form 3 independent experiments. **D-E**. Doxycycline-treated control and RAB5A-MCF10A cells were grown overlaid on top of Matrigel plugs for up to 21 days. Acini were fixed and processed for phase contrast imaging to monitor acini shape and size (left images) or, at various time point, for immunofluorescence to detect apoptotic caspase+ and proliferating, Ki67+ cells (see Fig. S4). Exemplar, phase contrast images are shown, (see also Movie S15). Scale Bar, 100 μm. In (**E)**, The average size of acini was quantified and expressed as mean±SD (n=100 acini/conditions in 5 independent experiments. *p<0.05, ** p<0.01. P value were calculated using each-pair Student’s t*-*test. **F**.Control and RAB5A-MCF10A cells were grown overlaid on top of Matrigel plugs for 14 days to allow full differentiation into hollow acini. Doxycycline was then added (time schedule of drug administration is on the top) to induce RAB5A expression in the presence or absence of PD0325901 and after 6 days acini were fixed and stained as indicated. Scale Bar, 80 μm. The size of acini was calculated by measuring their diameter. The number of KI67+ acini is also reported. Data are mean±SD (n=25 acini/conditions in 3 independent experiments). ** p<0.01. P value were calculated using each-pair Student’s t-test. **G**.Doxycycline-treated control and RAB5A-MCF10A cells were grown overlaid on top of mixed Matrigel:Collagene Type I (1:1) plugs for 21 days. Acini were fixed and processed for phase contrast imaging to monitor acini shape and size (left images). Exemplar phase contrast images are shown. Scale bar, 100 μm. Area of acini and acini roundness was quantified and expressed as mean±SD (n=40 acini/conditions in 5 independent experiments). ** p<0.01, paired Student’s t*-*test.

The ERK1/2-dependent re-awakening of collective motion and proliferation of terminal differentiated mammary glands has been associated with the initiation of a more complex program of branched morphogenesis that begins with the formation of multicellular buds^64, 65^. The latter process is thought to require in addition to specific growth factors, also the interaction of epithelial acini with the microenvironment and ECM components^66-68^. Collagen Type I, for example, has been used to increase mechanical tension and facilitate duct morphogenesis ^69, 70^. Henceforth, we grew MCF10A cells overlaid on mixed matrigel:collagen (5:1) gels^67, 71^. Under these conditions, cells form fully differentiated acini undistinguishable from those grown on Matrigel only (Fig. 6G). Addition of doxycycline to induce RAB5A expression, however, caused cysts to lose their spherical roundness, and promoted the formation of multicellular buds (Fig. 6G). Thus, RAB5A-dependent reawakening of cell motility occurs during 3D morphogenetic processes and enables to overcome proliferation arrest in fully differentiated epithelial cell assemblies.

### Endocytic unjamming promotes collective invasion in BC DCIS spheroids and in ex vivo BC foci

The discovery that unjamming impacts on the collective motility and on the growth dynamics of normal epithelial 3D ensembles prompted us to assess whether these processes can be exploited by BC to enhance their collective motility and invasive dynamic behavior^72^. To this end, we generated doxycycline-inducible MCF10.DCIS.com cells. These cells are isogenic to MCF10A, express oncogenic T24-H-RAS, a relative rare mutation in human BC lesions, but were derived from multiple-passage, murine orthotopic xenografts and shown to recapitulate in vivo and in vitro the progression from DCIS to invasive carcinoma^73^. During the DCIS phase, they grow under intra-ductal confinement where extreme cell packing and density exert mechanical stress, supress motility and tumour progression. Consistently, MCF10.DCIS.com cells are locked in jammed kinetically arrested state when plated at high confluency in 2D (Fig. S5A and ref.^31^). The expression of RAB5A promoted the reawakening of collective motion (Fig. S5A and Movie 16 and ref.^31^). It also accelerated directed motility of wounded monolayers, which instead of arresting after the opposing fronts collided, kept on flowing as collective streams, reminiscent of “a wound that never heals^74^” (Fig. 7A ad Movie 17). Biochemically, RAB5A expression decreased total EGFR levels, but robustly increased ERK1/2 without affecting AKT or p38 (which was slightly reduced) phosphorylation, as seen in untransformed MCF10A cells (Fig. S5B).

**Figure 7.**
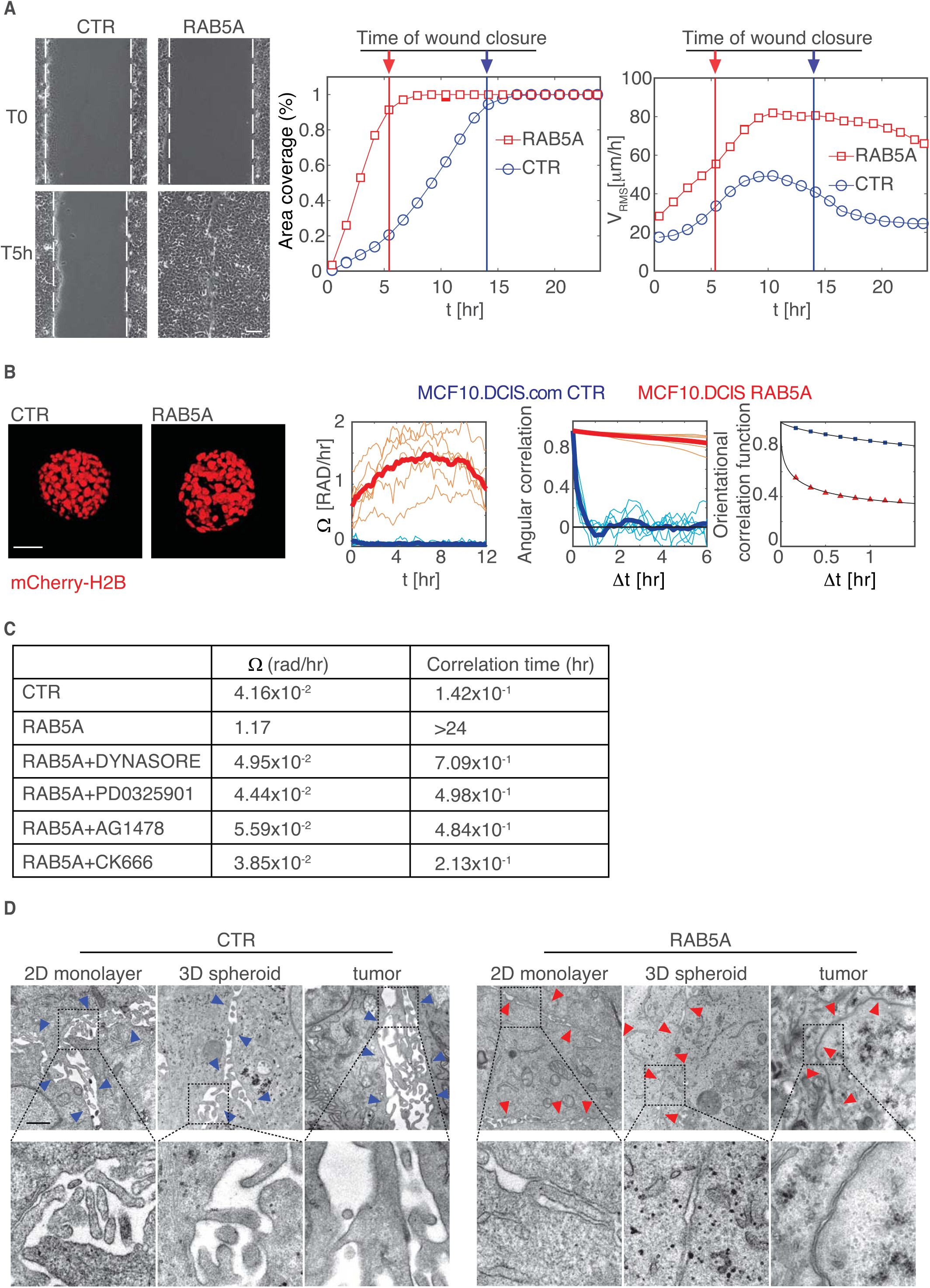
RAB5A-mediated unjamming promotes the emergence of coordinated angular rotation mode of breast cancer spheroids. **A**.Scratched wound migration of doxycycline-treated control and RAB5A-MCF10.DCIS.com seeded at jamming density. Representative still images at the indicated time points are shown. Dashed lines mark the wound edges. Scale bar, 100 μm. Motility was quantified by measuring (left) the percentage of area covered over time (calculations made with MATLABb software). On the most righthand, we also quantified *v*_*RMS*_ as function of time using PIV, along the entire movies to reveal that while control cells ceased migration, RAB5A-MCF10.DCIS.com keep on flowing (see Movie S17). Vertical bars indicate the time at which wounds close. Data are representative of 1 experiment out of >10 that were performed with similar outcome. **B**.Snapshots of control and RAB5A-MCF10.DCIS.com expressing mCherry-H2B that were grown as spheroids in low attachments plates. Spheroids were then embedded in thick 6.0 mg/ml of native Collagen Type I gels. After addition of doxycycline, the kinematic of spheroids was monitored by persistence of the rotational motion persistence of the rotational motion (Movie S18). A variance-based analysis was performed to extract the angular velocity Ω (expressed as rad/hr) of the spheroids as a function of time. The persistence of the rotational motion is quantified by considering the decay of the orientational correlation function, while the non-rigid part of the motion the spheroids, involving mutual cell rearrangement and fluid-like dynamics, is captured by the so-called overlap parameter *Q* (see Methods for details). The analysis was performed on 5-8 spheroid/conditions out of 3 independent experiments. **C**.inematic parameters of doxycycline-treated control and RAB5A-MCF10.DCIS.com expressing mCherry-H2B spheroids under the experimental conditions described above but treated with vehicle or the various indicated inhibitors (Movie S19). Mean angular velocity Ω and correlation time, extracted from an exponential fit of the orientational correlation functions, are reported (n=5 spheroids/condition out of 3 independent experiments). **D.**.Representative electron microscopy micrographs of doxycycline-treated control and RAB5A-MCF10.DCIS.com cells that were seeded at jamming density as 2D monolayers, allowed to form 3D spheroids which were embedded into thick, native Collagen type I gels, or injected into the mammary fat pad of immune-compromised mice. Blue arrows point to large spaces between cell–cell contacts, red arrows to tight cell–cell contacts. Scale bars, 2 μm.

In contrast with MCF10A, MCF10.DCIS.com cells form filled spheroids when grown in low attachment, or on top or embedded in 3D ECM matrix, and can generate invasive 3D outgrowth recapitulating DCIS-to-IDC conversion^75^. To test the impact of RAB5A-mediated unjamming on 3D growth dynamic and collective invasion of MCF10.DCIS.com, we generated mCherry-H2B control and RAB5A expressing cells, grew them as spheroids that were embedded in thick, native collagen type I to recapitulate the desmoplastic reactive environments of DCIS. After doxycycline addition, we monitored the spheroid kinematics. Whereas control cells display a slow, uncorrelated, disordered motion (Fig. 7B and Movie S18), RAB5A-MCF-10.DCIS.com cells acquired a striking CAM, displaying a remarkably large angular velocity Ω (of the order of ~12 rad/hr) and a strong persistence in the orientation of the instantaneous axis of rotation captured by the decay time of the orientational correlation function (the typical time interval over which the axis of rotation loses memory of its initial orientation), which is of the order of minutes in control, but > 24hr in RAB5A-MCF-10.DCIS.com spheroids (Fig. 7B). On top of this global rotational motion, a marked elevation of fluid-like motility of the cells is also observed. The characteristic time scale associated with this internal rearrangement dynamics is estimated by calculating the so-called overlap parameter *Q* (see Methods for details). Briefly, *Q* is a function of time that decays from 1 to 0 according to the number of nuclei that have been substantially displaced from their original position, when observed in a reference frame co-moving with the whole spheroid (Fig. 7B). The decay of *Q* does not depend on the rigid motion of the spheroid as a whole, but captures, instead, the “fluid-like” relative motion of the cells.

Endocytic reawakening of liquid-like CAM in 3D RAB5A-MCF-10.DCIS.com spheroids was dependent on EGFR activity, ERK1/2 phosphorylation, dynamin endocytosis and abrogated following inhibition of ARP2/3-mediated branched polymerization (Fig 7C-table and Movies S19). Furthermore, EM morphological analysis of control and RAB5A expressing monolayers, spheroids and orthotopically-injected tumours revealed that RAB5A induces junctional straightening and increases cell-cell surface contact areas (Fig. 7D). Thus, similar cellular/biochemical processes driving 2D locomotion and 3D acini morphogenesis operate in controlling the dynamic behaviour of oncogenic epithelial ensembles.

Next, we explored the pathological consequence of endocytic-mediated, unjamming of oncogenic collectives. Using mixed EGFP-LifeAct and mCherry-H2B-expressing spheroids embedded into collagen type I matrix, we monitored their behaviour over longer time scale. We invariably observed that RAB5A promoted CAM, followed by the extension into the surrounding ECM of multicellular protrusion in the form of invasive buds and strands, suggesting that unjamming and collective invasion might be temporally coordinated and possibly coupled (Fig. 8A and Movie S20).

**Figure 8.**
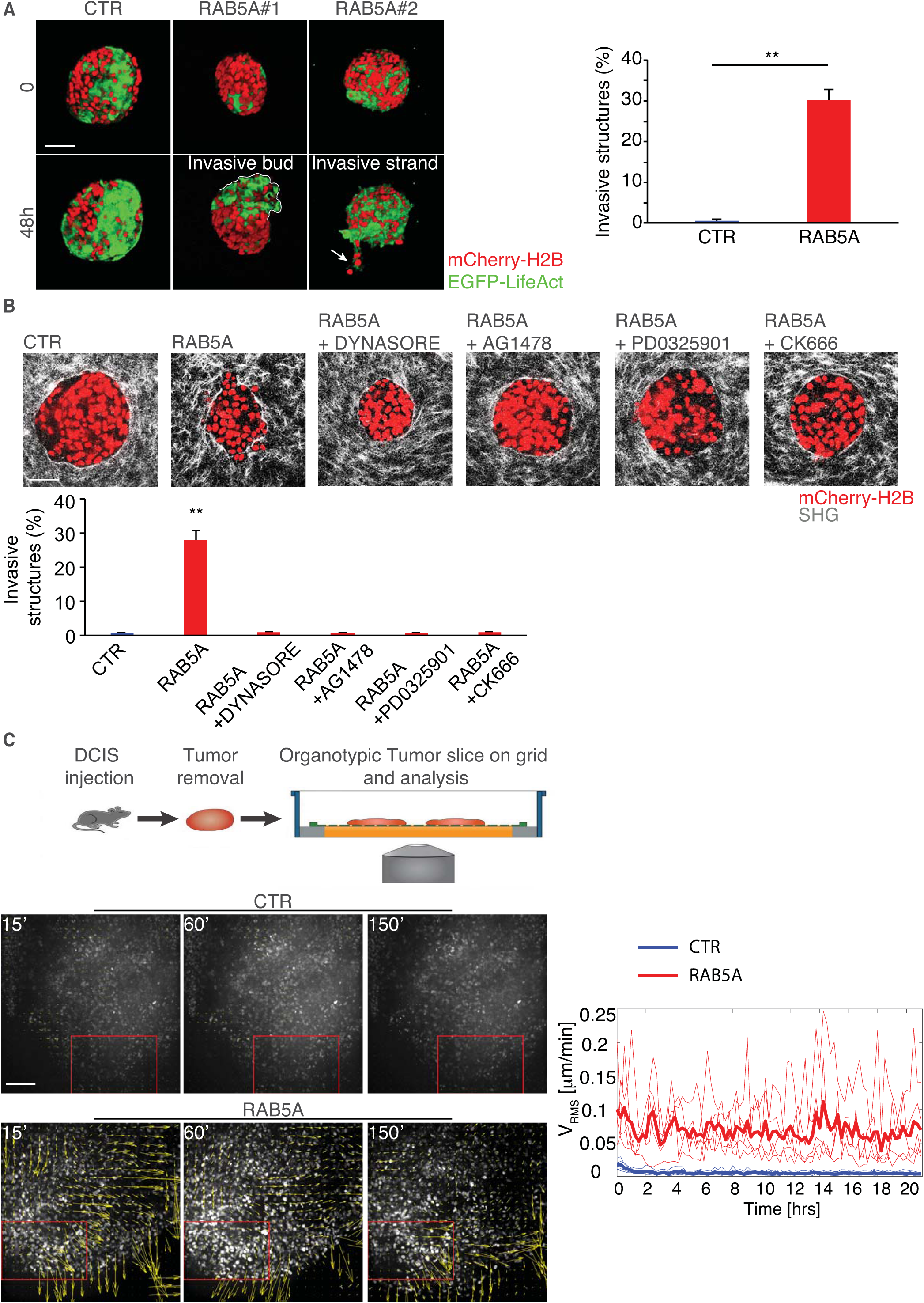
RAB5A-mediated 3D unjamming promotes collective invasion in tumour spheroids and in ex vivo DCIS tumours slices. **A**.Control and RAB5A-MCF10.DCIS.com expressing mCherry-H2B and EGFP-LifeAct were grown as spheroid in low attachment. Spheroids were then embedded in thick 6.0 mg/ml of native Collagen Type I gels. After addition of doxycycline, the kinematic of spheroids and the formation of invasive multicellular structures was monitored by confocal time lapse microscopy and snapshots are shown (Movie S20). The line delineates an invasive multicellular bud; the arrow points to an invasive strand. The percentage of spheroids with invasive multicellular structures was scored and expressed as mean±SD (n=15/experimental conditions in 5 independent experiments). ** p<0.01, calculated using Student’s t-test. Scale Bar, 150 μm. **B**.Analysis of Collagen type I structures using SHG of doxycycline-treated Control and RAB5A-MCF10.DCIS.com expressing mCherry-H2B spheroids embedded into 6.0 mg/ml of native Collagen Type I gels in the presence of vehicle or the indicated inhibitors. The percentage of spheroids with invasive multicellular structures was scored and expressed as mean±SD (n=15/experimental conditions in 5 independent experiments). ** p<0.01, calculated using each-pair Student’s t-test. Scale Bar, 70 μm. **C**.Schematic of the experimental design. Control and RAB5A-MCF10.DCIS.com expressing mCherry-H2B and EGFP-LifeAct cells (2X10^5^) were injected into the mammary fat pad of immunocompromised mice. After 4 weeks, tumours that forms ductal carcinoma in situ lesion that are progression to become IDC were mechanically excised and tumour tissues slices were placed on a grid at the air-liquid interface. The organotypic cultures were treated with doxycycline and monitored by time-lapse confocal microscopy (Movie S21) for 24hr. Snapshots of the velocity fields obtained from PIV analysis of motion (Movie 22), which was used to extract the root mean square velocity, *v*_*RMS*_, as function of time (Right plot). Boxed areas indicate representative fields of view used for the analysis (At least 5 field of view/movies of 3 independent experiments were analysed). Thin lines in the right plot indicate the actual evolution of *v*_*RMS*_/ in each of the field of view, while thick lines are the average of the *v*_*RMS*_. Scale Bar, 150 μm.

Collective invasion into native collagen type I, which, at the concentration used, form a very dense network of intricate fibres, can only occur following its remodelling. Consistently, Second Harmonic Generation (SHG) signals of two-photon illumination revealed that RAB5A-expressing spheroids extensively remodelled the fibrillary collagen, generating gaps and channels for collective invasion to occur (Fig. S5C and Fig. 8B). EGFR, ERK1/2, Dynasore, and ARP2/3 inhibitor of CAM prevented also collagen remodelling and the formation of invasive buds, suggesting that acquisition of unjamming in 3D promote collective invasion (Fig. 8B).

Finally, we tested the latter hypothesis in a closer to physio-pathological conditions using *ex vivo* organotypic tumour slices from DCIS injected into the mammary fat pad of immunocompromised mice. To this end, m-Cherry-H2B and EGFP-LifeAct-expressing-control and RAB5A-MCF10.DCIS.com were xenotransplanted into NSG mice. After 4 weeks, tumour masses were mechanically excised, and organotypic tissue slices were grown at the air-liquid interface (see methods and ref.^76^). Under such conditions, bulk tumour tissues remain viable for up to 2 weeks and, more importantly, their dynamic behaviour can be tracked over hours with high temporal resolution. Control and RAB5A-DCIS organotypic cultures were exposed to doxycycline to induce transgene expression and monitored by time lapse confocal microscopy. Whereas control tumours were largely immobile (Fig. 8C and Movies S21), jammed and compacted, the expression of RAB5A induces reawakening of cell dynamics. Cells became highly motile and appeared to stream in a way that resembles currents in a river (Fig 8C and Movies S21). PIV analysis captured quantitatively the transition to collective locomotory mode (Movie S22).

We concluded that endocytic unjamming of kinetically arrested dense DCIS tumours is sufficient to instigate motility and to promote collective invasive behaviour.

## Discussion

Here, we provide the first evidence for a novel molecular route to unjamming, which reinstates the possibility of multicellular rearrangements in otherwise immobile mature epithelia and densely-packed carcinoma. Biochemically, we showed that elevated NCE internalization of EGFR promotes its accumulation into endosomal vesicles, which become proficient signalling platforms for the prolonged and elevated activation of ERK1/2. Our data using an endosomal-FRET-ERK1/2 sensor is, to our knowledge, the first direct demonstration of the latter contention. These findings are also consistent with the notion that while ERK1/2 signalling initiates at the plasma membranes, it continues into endosomes impacting not only on signal intensity and duration but also on specificity^46-49,77^. Accordingly, RAB5A-induced EGFR endosomal signalling promotes the hyper-phosphorylation of WAVE2 that, by controlling branched actin polymerization, contributes to the extension of oriented cryptic lamellipodia^60^. Physically, the latter protrusions exert increased traction forces^31, 78^, and enhance cell orientation, which is found to be the fundamental ingredient to obtain liquid states with large Vičsek-like polar alignment, a signature of flocking in jammed epithelia^31-33^.

It is likely that additional, not yet identified substrates are phosphorylated by endosomal ERK1/2. Indeed, reawakening of motility in jammed epithelia requires perturbations of different cellular and supra-cellular pathways and properties (including cell-cell adhesion, surface tension^79^ and monolayer rigidity^33^ for optimal long-range force transmission^80^, volume and density fluctuations^32, 81^), which are all required to initiate a mode of locomotion that combines large scale correlation length with increased local cell arrangement typical of fluidized RAB5A tissues. This notwithstanding, impairing protrusion extension impedes the emergence of persistent flocking motion and abrogates the re-awakening of motility, pointing to a major role of the protrusion extension mechanism in setting the local directionality of the flocking motion.

Our results also represent a step forward in addressing the physio-pathological relevance of tissue *unjamming*. In 3D morphogenetic assays of mammary gland morphogenesis, we showed that endo-ERK1/2-mediated unjamming not only promotes the acquisition of coordinated angular rotational mode of motion, but that it is sufficient to overcome differentiation-induced proliferation arrest. These latter two features combine with input arising from the presence of collagen type I in the substrate, which likely provides increased mechanical tension^69-71, 82^, to facilitate multicellular bud formation, a process marking the beginning of branching morphogenesis^68^. Indeed, local elevation of ERK1/2, promoting both re-entry into proliferation and collective motion, has been shown to mediate murine mammary gland branching morphogenesis^64, 65^, remarkably similar to what we observe in our in vitro 3D assays. Thus, we argue that the endocytic-dependent jamming transition molecularly described here might be a valuable framework to account for the initiation of complex morphogenetic processes.

Finally, we showed that Endo-ERK1/2-unjamming might also be sufficient to overcome the rigid, kinetically-silent state of packed epithelial carcinoma spheroids that grow confined and encased by a thick collagen type I matrix. Endocytic unjamming, here, leads to the acquisition of flocking fluid modes of motion, in which highly coordinated and collective rotational migration and local unjamming coexist, providing the first evidence that this transition occurs not only in 2D monolayers, but also in complex 3D environments. Re-awakening of collective motion, under these conditions, is accompanied by a dramatic remodelling of the ECM and by the extension of collective invasive buds and strands. This process recapitulates some aspects of the transition from DCIS, which grow under intra-ductal confinement where extreme cell packing and density exert mechanical stress, supress motility and tumour progression, to invasive carcinoma, which disperse locally also through collective invasion^83^. We showed that RAB5A is elevated in various human breast cancer subtypes and its elevated expression correlates with reduced relapse free probability, supporting the notion that jamming transition might, indeed, be an additional mechanism to promote local collective invasion of densely packed breast carcinoma. In this setting, RAB5A induction could be an alternate route to EGFR addiction in invasive breast carcinomas, independently of HER2 status.

## Methods

Methods and any associated accession codes and references are included after the references.

## Acknowledgments

This work has been supported by: the Associazione Italiana per la Ricerca sul Cancro (AIRC) to GS (IG#18621), PPDF (IG#18988 and MCO 10.000); the Italian Ministry of University and Scientific Research (MIUR) to PPDF; the Italian Ministry of Health (RF-2013-02358446) to GS. Regione Lombardia and CARIPLO foundation (Project 2016-0998) to RC; Worldwide Cancer Research (WCR#16-1245) to SS. CM and FG are partially supported by fellowships from the University of Milan, EB from the FIRC-AIRC.

## Author contributions

AP, CM, EF design and perform all the experiments and edited the manuscript, SC aid in generating cell lines and in the analysis of IF and kinematic studies, SB, SS and PPFD conceived internalization assays and interpreted trafficking results, GVB perform EM studies, EM, MG, and DP aided in all the imaging acquisition, FRET and PIV analysis, CT aided in analysis of RAB5A expression in BC, FG and RC analyzed all the kinematic data and developed pipeline for the analysis of 3D motility, edited the manuscript and conceived part of the study together with CM, GS conceived the whole study, wrote the manuscript and supervised the whole work.

## Competing financial interests

The authors declare no competing financial interests.

## Data Availability Statement

Codes used for the analysis are all indicated in the methods section. The authors declare that all data supporting the findings of this study are available within the paper and its supplementary information files and there no restriction on data availability.

## METHODS

### Plasmids, antibodies and reagents

Doxycycline-inducible lentiviral vectors pSLIK-neomycin (neo) carrying RAB5A or RAB5C sequences and pSLIK-hygromycin (hygro) carrying RAB5B sequence were obtained by Gataway Technology (Invitrogen), following the manufacturer’s protocol. The plasmids pBABE-puromycin (puro)-mCHERRY-H2B and pBABE-puro-EGFP-H2B were provided by IFOM-Imaging Facility. The lentiviral expression construct pRRL-Lifeact-EGFP-puromicin (puro) was a gift of Olivier Pertz (University of Basel, Basel, Switzerland). pBabe-Puro-MEK-S218D/S222D (MEK-DD) vector was purchased from Addgene.

FRET EKAREV-ERK1/2 sensor^1^ was generated by cloning synthetized FYVE domain of SARA into the BamHI/EcoRI cleaved EKAREV-FRET vector to generate pPBbsr2-3560NES-EKAREV-FRET new vector.

Mouse monoclonal antibodies raised against α-tubulin (#T5168) or vinculin (#V9131) were from Sigma-Aldrich. Rabbit polyclonal anti-RAB5A (S-19, #sc-309) and goat polyclonal anti-EEA-1 (N-19, #sc-6415) antibodies from Santa Cruz Biotechnology. Monoclonal rabbit anti-human RAB5A - ab109534, dilution 1:100, (Abcam[EPR5438]) was used of IHC;Rabbit polyclonal anti-Giantin (#PRB-114C) antibody was from Covance. Mouse monoclonal anti-human Ki-67 Antigen (MIB-1, #M7240) antibody was from Dako. Mouse monoclonal anti-AP50 (AP2mu) (31/AP50, #611350) was from BD Bioscience. Mouse monoclonal anti-E-cadherin (#610181) antibody was from Transduction Lab. Rabbit polyclonal anti-phospho-EGFR (Tyr1086, #2220), rabbit monoclonal anti-phospho-p44/42 MAPK (ERK1/2) (Thr202/Tyr204, #4370), rabbit polyclonal anti-p44/42 MAPK (ERK1/2) (#9102), rabbit monoclonal anti-phospho-p38 MAPK (Thr180/Tyr182, 3D7, #9215), mouse monoclonal anti-p38 MAPK (L53F8, #9228), rabbit monoclonal anti-phospho-AKT (Ser473, 193H12, #4058), rabbit polyclonal anti-AKT (#9272), rabbit polyclonal anti-MEK1/2 (#9122) and rabbit polyclonal anti-cleaved Caspase-3 (Asp175, #9661) antibodies were from Cell Signalling Technology. Rabbit polyclonal anti-phospho-WAVE2 (Ser343, #07-1512), rabbit polyclonal anti-phospho-WAVE2 (Ser351, #07-1514) and mouse monoclonal anti-Laminin-V (P3H9-2, #MAB1947) antibodies were from Merck/Millipore. Mouse monoclonal anti-WAVE2 antibody was homemade. Rabbit polyclonal anti EGFR (806), directed against aa 1172-1186 of human EGFR (ImmunoBlot) and mouse monoclonal anti-EGFR (m108 hybridoma) directed against the extracellular domain of human EGFR (IF) were a gift from P.P. Di Fiore. Secondary antibodies conjugated to horseradish peroxidase were from Bio-Rad (#7074, #7076); Cy3-secondary antibodies from Jackson ImmunoResearch (#711-165-152, #715-165-150); DAPI (#D-1306) and AlexaFluor 488 (A-11055, A-21202) were from Thermo Fisher Scientific. TRITC-(#P1951) and FITC-(#P5282) conjugated phalloidin were from Sigma Aldrich.

Doxycycline Hyclate (DOX, #D9891), Dynasore Hydrate (#D7693), AG1478 (#T4182) and CK666 (#SML0006) were from Sigma Aldrich. PD0325901 (#444966) was from Merck/Millipore.

### Cell cultures and transfection

MCF10A cells were a kind gift of J. S. Brugge (Department of Cell Biology, Harvard Medical School, Boston, USA) and were maintained in Dulbecco’s Modified Eagle Medium: Nutrient Mixture F-12 (DMEM/F12) medium (Biowest) supplemented with 5% horse serum, 1% L-Glutamine (EuroClone), 0.5 mg ml^-1^hydrocortisone (Sigma-Aldrich), 100 ng ml^-1^ cholera toxin (Sigma-Aldrich), 10 µg ml^-1^ insulin (Sigma-Aldrich) and 20 ng ml^-1^ EGF (Vinci Biochem). MCF10.DCIS.com cells were kindly provided by J. F. Marshall (Barts Cancer Institute, Queen Mary University of London, UK) and maintained in Dulbecco’s Modified Eagle Medium: Nutrient Mixture F-12 (DMEM/F12) medium supplemented with 5% horse serum, 1% L-Glutamine, 0.5 mg ml^-1^hydrocortisone, 10 µg ml^-1^ insulin and 20 ng ml^-1^ EGF. All cell lines have been authenticated by cell fingerprinting and tested for mycoplasma contamination. Cells were grown at 37 °C in humidified atmosphere with 5% CO2. MCF10A cells were infected with pSLIK-neo-EV (empty vector-CTR), pSLIK-neo-RAB5A, pSLIK-hygro-RAB5B or pSLIK-neo-RAB5C lentiviruses and selected with the appropriate antibiotic to obtain stable inducible cell lines. MCF10.DCIS.com were infected with pSLIK-neo-EV (empty vector-CTR) or pSLIK-neo-RAB5A lentiviruses and selected with the appropriate antibiotic to obtain stable inducible cell lines. Constitutive expression of EGFP-LifeAct- or mCHERRY- or EGFP-H2B was achieved by lentiviral and retroviral infection of MCF10A and MCF10DCIS.com cells with EGFP-LifeAct-puro or pBABE-puro-mCHERRY-H2B/ pBABE-puro-EGFP-H2B vectors, respectively.

Transfections were performed using either calcium phosphate or FUGENE HD Transfection reagent (#E2311, PROMEGA) reagents, according to manufacturer’s instructions. FUGENE HD reagent was used for FRET-EKAREV-ERK1/2 transfection in MCF10A cells.

### Generation of lentiviral and retroviral particles

Packaging of lentiviruses or retroviruses was performed following standard protocols. Viral supernatants were collected and filtered through 0.45 µm filters. Cells were subjected to four cycles of infection and selected using the appropriate antibiotic: neomycin for pSLIK-neo vector (150µg/ml), hygromycin for pSLIK-hygro vector (100 µg/ml) or puromycin for EGFP-LifeAct or pBABE vectors (2 µg/ml). After several passages, stable bulk populations were selected and induced by Doxycycline Hyclate (2.5 µg/ml) in order to test: i) induction efficiency by Western Blotting and quantitative RT-PCR (qRT-PCR), and ii) the homogeneity of the cell pool by immunofluorescence staining, as previously shown^2^.

### RNA interference

siRNAs (small interfering RNAs) delivery was achieved by mixing 1 nM of specific siRNAs with Optimem and Lipofectamine RNAiMAX Transfection Reagent (Life Technologies). The first cycle of interference (reverse transfection) was performed on cells in suspension. The day after, a second cycle of interference (forward transfection) was performed on cells in adhesion. The following siRNAs were used for knocking down specific genes. All sequences are 5’ to 3’.

Dynamin2 (DNM2): 5’-GACATGATCCTGCAGTTCA-3’ (Dharmacon) AP2mu: 5’-UGACCCGAAAGGCAUCCACCCCC-3’ (Riboxx)

MP1 (LAMTOR3): 5’-CAAUUUAAUCGUUUACCUU-3’ (Silencer Select, Ambion)

P14 (LAMTOR2): 5’-CCCAAGUGGCGGCAUCUUA-3’ (Silencer Select, Ambion)

Reticulon 3 (RTN3): 5’-CCCUGAAACUCAUUAUUCGUCUCUU-3’ (Stealth, Invitrogen)

Reticulon 4 (RTN4): 5’-CCAGCCUAUUCCUGCUGCUUUCAUU-3’ (Stealth, Invitrogen)

For each RNA interference experiment, negative control was performed with the same amount of scrambled siRNAs. Silencing efficiency was controlled by qRT-PCR.

### Quantitative RT-PCR analysis

Quantitative RT-PCR analysis was performed as previously shown^2^. Total RNA was extracted using RNeasy Mini kit (Qiagen) and quantified by NanoDrop to assess both concentration and quality of the samples. Reverse transcription was performed using SuperScript VILO cDNA Synthesis kit from Invitrogen. Gene expression was analyzed using TaqMan Gene expression Assay (Applied Biosystems). 0.1 ng of cDNA was amplified, in triplicate, in a reaction volume of 25 µl with 10 pMol of each gene-specific primer and the SYBR-green PCR MasterMix (Applied Biosystems). Real-time PCR was performed on the 14 ABI/Prism 7700 Sequence Detector System (PerkinElmer/Applied Biosystems), using a pre-PCR step of 10 min at 95°C, followed by 40 cycles of 15 s at 95°C and 60 s at 60°C. Specificity of the amplified products was confirmed by melting curve analysis (Dissociation Curve TM; PerkinElmer/Applied Biosystems) and by 6% PAGE. Preparations with RNA template without reverse transcription were used as negative controls. Samples were amplified with primers for each gene (for details see the Q-PCR primer list below) and GAPDH as a housekeeping gene. The Ct values were normalized to the GAPDH curve. PCR experiments were performed in triplicate and standard deviations calculated and displayed as error bars. Primer assay IDs were: GAPDH, Hs99999905_m1; RAB5A, Hs00702360_s1; RAB5B, Hs00161184_m1 and RAB5C, Hs00428044_m1, Dynamin2 (DNM2) Hs00974698_m1, MP1 (LAMTOR3) Hs00179753_m1, P14 (LAMTOR2) Hs00203981_m1, Reticulon3 (RTN3) Hs01581965_m1, Reticulon4 (RTN4) Hs01103689_m1.

### Immunoblotting

For protein extraction, cells, previously washed with cold PBS, were lysed in JS buffer supplemented with proteases and phosphatases inhibitors [50 mM HEPES PH 7.5, 50 mM NaCl, 1% glycerol; 1% Triton X-100, 1.5 mM MgCl_2_. 5 mM EGTA plus protease inhibitor cocktail (Roche, Basel, Switzerland), 1 mM DTT, 20 mM Na pyrophosphate pH 7.5, 50 mM NaF, 0.5 M Na-vanadate in HEPES pH 7.5 to inhibit phosphatases]. Lysates were incubated on ice for 10 minutes and cleared by centrifugation at 13,000 rpm for 30 min at 4°C. Protein concentration was quantified by Bradford colorimetric protein assay. The same amount of protein lysates was loaded onto polyacrylamide gel in 5X SDS sample buffer. Proteins were transferred onto Protran Nitrocellulose Transfer membrane (Whatman), probed with the appropriate antibodies and visualized with ECL western blotting detection reagents (GE Healthcare). Membrane blocking and incubation in primary or secondary antibodies were performed for 1h in TBS/0.1% Tween/5% milk for antibodies recognizing the total proteins or in TBS/0.1% Tween/5% BSA for antibodies recognizing phosphorylated proteins.

### Immunohistochemistry on DCIS and IDC

Sections from archival human breast cancer samples were collected from the archives of the Tumor Immunology Laboratory of the Human Pathology Section, Department of Health Sciences, University of Palermo, Italy.

Immunohistochemistry was performed using a polymer detection method (Novolink Polymer Detection Systems Novocastra, Leica Biosystems, Newcastle, Product No: RE7280-K). Tissue samples were fixed in 10% buffered formalin and embedded in paraffin. Four-micrometers-thick tissue sections were dewaxed and rehydrated. The antigen unmasking technique was performed using Novocastra Epitope Retrieval Solution pH6 citrate-based buffer in thermostatic water bath at 98°C for 30 minutes. Subsequently, the sections were brought to room temperature and washed in PBS-Tween. After neutralization of the endogenous peroxidase with 3% H_2_O_2_ and protein blocking by a specific protein block, the samples were incubated 1h with monoclonal rabbit anti-human RAB5A [EPR5438] - ab109534 (dilution 1:100, Abcam). Staining was revealed by polymer detection kit (Novocastra, Ltd) and AEC (3-Amino-9-Ethylcarbazole) substrate chromogen. The slides were counterstained with Harris hematoxylin (Novocastra, Ltd). All the sections were analyzed under a Zeiss Axio Scope A1 optical microscope (Zeiss, Germany) and microphotographs were collected using an Axiocam 503 Color digital camera with the ZEN2 imaging software (Zeiss Germany)

### Cell streaming and wound healing assays

As previously shown^2^, cells were seeded in 6-well plate (1.5*10^6^ cells/well) in complete medium and cultured until a uniform monolayer had formed. RAB5A expression was induced, were indicated, 16 hours before performing the experiment by adding fresh complete media supplemented with 2.5 µg/ml Doxycycline Hyclate to cells. Comparable cell confluence was tested by taking pictures by differential interference contrast (DIC) imaging using a 10x objective and counting the number of nuclei/field. In cell streaming assay, medium has been refreshed before starting imaging. In wound healing assay, cells monolayer was scratched with a pipette tip and carefully washed with 1X PBS to remove floating cells and create a cell-free wound area. The closure of the wound was monitored by time-lapse. Olympus ScanR inverted microscope with 10x objective was used to take pictures every 5-10 minutes over a 24 hours period (as indicated in the figure legends). The assay was performed using an environmental microscope incubator set to 37°C and 5% CO2 perfusion. After cell induction, Doxycycline Hyclate was maintained in the media for the total duration of the time-lapse experiment. The percentage of area covered by cells (area coverage %) overtime and wound front speed were calculated by MatLab software. In chemical inhibitors experiments, the inhibitor was added together with Doxycycline Hyclate in fresh media 1 h before starting imaging. For cell streaming assay performed on interfered cells, cells were interfered in suspension (first cycle) and directly plated at the desired concentration, following the same conditions already described in “RNA interference” section.

For detection of cryptic lamellipodia, MCF10A cells stably expressing EGFP-LifeAct were mixed in a 1:10 ratio with unlabeled cells and seeded in cell streaming assay, as described before. Cell migration was monitored by time-lapse phase contrast and fluorescence microscopy, collecting images at multiple stage positions in each time loop. Olympus ScanR inverted microscope with 20x objective (+1.6x Optovar) was used to take pictures every 90 seconds. Each assay was done 5 times and at least 25 cells/condition were counted in each experiment. Where indicated, PD0325901 was added 1 h before imaging.

### FRET Analysis

Using a customised macro in ImageJ, FRET data were analysed using the ratiometric approach. CFP, YFP and FRET images were background subtracted, converted in 32bits and the smoothed YFP image were tresholded and used as a mask to highlight the vesicular-like structures of interest. On these areas the average FRET/CFP ratio was then calculated as described in Kardash E. et al.^3^

### 3D morphogenesis assay

MCF10A morphogenesis assay was per formed as already described^67^. Briefly, MCF10A cells were trypsinized and resuspended in MCF10A culture medium. Eight-well chamber slides (#80826 IBIDI) were coated with 40 μl/well of Growth Factor Reduced Matrigel Matrix Basement Membrane HC 10.2 mg/ml (#354263, Corning) or with 1:1 mixture of Matrigel HC 10.2 mg/ml and Type I Bovine Collagen 3 mg/ml (#5005 Advanced BioMatrix). Once the matrix is polymerized, 2.5*10^3^ cells were plated into each well on the top of the matrix layer in culture medium supplemented with 2% Matrigel HC 10.2 mg/ml and 5 ng/ml EGF. Complete acini morphogenesis was allowed by incubating the cells for 3 weeks and replacing assay media every four days.

On day 21 acini were treated with 2.5 μg/ml Doxycycline Hyclate to induce RAB5A expression. Cells were maintained under stimulation for 6 days, changing the medium every 2 days, before fixation with 4% paraformaldehyde (PFA) and stained with specific antibodies. When inhibitors were used, the media were refreshed every day.

### 3D spheroid kinematic assay

MCF10DCIS.com cells were plated on Ultra-Low attachment surface 6-well plate (#3471 CORNING) at a density of 5*10^3^ cells/well. Cells were grown in serum-free condition for 10 days by adding fresh culture media every 2 days. Then every single well of spheres were collected and resuspended in 150 μl of 6 mg/ml Collagen Type I (#35429 CORNING), diluted in culture media, 50 mM Hepes, 0,12 NaHCO3 and 5 mM NaOH. The unpolymerized mix sphere/collagen was placed in Eight-well chamber slides and incubated at 37°C for o/n. The day after, before imaging, 2.5μg/ml Doxycycline Hyclate was added over the polymerized collagen mix to induce RAB5A expression.

### Ex Vivo DCIS tumor slice motility assay

All animal experiments were approved by the OPBA (Organisms for the well-being of the animal) of IFOM and Cogentech. All experiments complied with national guidelines and legislation for animal experimentation. All mice were bred and maintained under specific pathogen-free conditions in our animal facilities at Cogentech Consortium at the FIRC Institute of Molecular Oncology Foundation and at the European Institute of Oncology in Milan, under the authorization from the Italian Ministry of Health (Autorizzazione N° 604-2016).

For mammary fat pad tumor development in NSG mice MCF10DCIS.com cells were trypsin detached, washed twice, and resuspended in PBS to a final concentration 2*10^5^/13 µl. The cell suspension was then mixed with 5 µl growth factor–reduced Matrigel and 2 µl Trypan blue solution and maintained on ice until injection. Aseptic conditions under a laminar flow hood were used throughout the surgical procedure. Female NOD.Cg-PrkdcscidIl2rgtm1Wjl/SzJ (commonly known as the NOD SCID gamma; NSG) mice, 6–9 weeks old, were anesthetized with 375 mg/Kg Avertin, laid on their backs, and injected with a 20-µl cell suspension directly in the fourth mammary fad pad. After 4 weeks mice were sacrificed and the primary tumors were removed, cut by a scalpel and each tumor slide was placed over a metal grid inserted in 6-well plate to allow tumors to grow on an interface air/culture medium. Before imaging, 2.5μg/ml Doxycycline Hyclate was added to tumor slices culture media to induce RAB5A expression. Tumor cells were maintained under stimulation for 3 days, changing the medium every day.

### Immunofluorescence

As previously shown^2^, cells were fixed in 4% paraformaldehyde (PFA) and permeabilized with 0.1% Triton X-100 and 1% BSA 10 minutes (except for EEA-1 staining, permeabilized with 0.02% Saponin and 1% BSA 10 minutes and pERK1/2 staining, permeabilized with ice cold 100% Methanol for 10 minutes). In EGFR staining experiments, permeabilization step was avoided where indicated (non-permeabilized conditions) in order to detect only total cell surface EGFR. After 1X PBS wash, primary antibodies were added for 1 hour at room temperature. Coverslips were washed in 1X PBS before secondary antibody incubation 1 hour at room temperature, protected from light. FITC- or TRITC-phalloidin was added in the secondary antibody step, where applicable. After removal of not specifically bound antibodies by 1X PBS washing, nuclei were stained with 0.5 ng/ml DAPI. Samples were post-fixed and mounted on glass slides in anti-fade mounting medium (Mowiol). Antibodies were diluted in 1X PBS and 1% BSA. Images were acquired by wide-field fluorescence microscope or confocal microscope, as indicated in figure legends.

Immunofluorescence on MCF10A-derived acini was performed by fixing acini with 4% paraformaldehyde for 20 minutes at RT. Then cells were permeabilized with 0.5% TRITON X-100 in PBS for 10 minutes at 4°C and incubated with blocking solution (PBS + 0.1% BSA + 10% goat serum) for 1 hour at RT. Acini were incubated with indicated primary antibodies diluted in blocking solution for o/n at 4°C. The day after acini were incubated with indicated secondary antibodies diluted in blocking solution for 1 hour at RT. Finally, acini were incubated with DAPI in PBS for 20 minutes at RT. Samples were then maintained at 4°C in PBS before imaging.

E-cadherin staining was analysed by confocal microscopy and images were processed to obtain the straightness index of the junction. “Junction length” was measured by tracking a straight line and “junction tracking” was obtained by tracking manually the same junction following its profile. The straightness index of the junction has been quantified as the ratio of the junction length and the junction tracking.

### ^125^I-EGF internalization assay

Internalization of ^125^I-EGF was performed at low EGF (1 ng/ml) or high EGF (30 ng/ml) as described in ref.^4^.

Briefly, MCF10A cells were plated in 24-well plates in at least duplicate for each time point, plus one well to assess non-specific binding. Cell monolayers were EGF-starved 24 hours and induced overnight by Doxycycline Hyclate. The day after cells were incubated in assay medium (DMEM/F12 supplemented with Cholera Toxin (100 ng/ml), 0,1% BSA, 20mM Hepes, DOX (2.5µg/ml) and then incubated at 37°C in the presence of 1 ng/ml ^125^I-EGF, or 30 ng/ml EGF (1 ng/ml ^125^I-EGF (Perkin Elmer) + 29 ng/ml cold EGF. At different time points (2, 4, 6 min) the amount of bound ^125^I-EGF was measured with an acid wash solution pH 2.5 (0.2 M acetic acid, 0.5 M NaCl). Cells were then lysed with 1N NaOH, which represents the amount of internalized ^125^I-EGF. Non-specific binding was measured at each time point in the presence of an excess of non-radioactive EGF (300 times). After being corrected for non-specific binding, the rate of internalisation was calculated as the ratio between internalised and surface-bound radioactivity. Surface EGFRs were measured by ^125^I-EGF saturation binding as described^5^.

### EGF recycling assay

Recycling assays of ^125^I-EGF were performed as described in^5^. In brief, cell monolayers were EGF-starved 24 hours and induced overnight by Doxycycline Hyclate. The day after cells were incubated in assay medium (DMEM/F12 supplemented with Cholera Toxin (100ng/ml), 0,1% BSA, 20mM Hepes, DOX (2.5µg/ml), then incubated with ^125^I-EGF (30 ng/ml: 5 ng/ml of ^125^I-EGF + 25 ng/ml of cold EGF) for 15 min at 37 °C, followed by mild acid/salt treatment (buffer at pH 4.5, 0.2 M Na acetate pH 4.5, 0.5 M NaCl) to remove bound EGF. Cells were then chased at 37°C in a medium containing 4 µg/ml EGF for the indicated times, to allow internalization and recycling. At the end of each chase time, the medium was collected, half was counted directly (free) and half was subjected to TCA precipitation to determine the amount of intact/recycled (TCA precipitable) and degraded (TCA soluble) ^125^I-EGF present in it. Surface-bound ^125^I-EGF was extracted by acid treatment (0.5M NaCl, 0.2M acid acetic). Finally, cells were lysed in 1N NaOH to determine intracellular ^125^I-EGF. Data are expressed as the fraction of intact ^125^I-EGF in the medium with respect to the total (total medium + total surface + total intracellular). Non-specific counts were measured for each time point in the presence of a 300-fold excess of cold ligand, and were never >3-10 % of the total counts.

### Image acquisition

Time-lapse imaging of 3D acini/spheroids motility was performed on a Leica TCS SP8 laser confocal scanner mounted on a Leica DMi8 microscope equipped with motorized stage; a HC PL FLUOTAR 20X/0.5NA dry objective was used. A white light laser was used as illumination source. LAS X was the software used for all the acquisitions.

Image acquisition conditions were set to remove channel crosstalk, optimizing spectral detection bands and scanning modalities. ImageJ software was used for data analysis.

Collagen SHG analysis on collagen embedded MCF10DCIS spheroids was performed with a confocal microscope (Leica; TCS SP5) on an upright microscope (DM6000 CFS) equipped with blue (argon, 488 nm), yellow (561 nm solid state laser), and red (633 nm solid state laser) excitation laser lines with an HCX PL APO 40X/1.25-0.75NA oil immersion objective and controlled by Leica LAS AF software (Leica). We used a two-photon excitation (2PE) technique with a pulsed infrared laser (Chameleon Ultra II; Coherent) at 980 nm.

EKAREV FRET analysis was performed using a DeltaVision Elite imaging system (Applied Precision) controlled by softWoRx Explorer 2.0 (Applied Precision) equipped with a DV Elite CMOS camera and an inverted microscope (IX71; Olympus) using a PlanApo N 60X/1.42NA oil-immersion objective lens.

Ex vivo DCIS tumor slice motility assay was performed using an Olympus IX83 inverted microscope controlled by CellSens software (Olympus) and equipped with a iXon Ultra Andor (EMCCD) 16 bit camera using a UplanSApo 10X/0.4NA dry objective.

### Electron Microscopy

Electron microscopic examination was performed as previously described^6, 7^. A description of each process is described below.

Embedding: the tissue and 3D spheroids were fixed with of 4% paraformaldehyde and 2.5% glutaraldehyde (EMS, USA) mixture in 0.2 M sodium cacodylate pH 7.2 for 2 hours at RT, followed by 6 washes in 0.2 sodium cacodylate pH 7.2 at RT. Then cells were incubated in 1:1 mixture of 2% osmium tetraoxide and 3% potassium ferrocyanide for 1 hour at RT followed by 6 times rinsing in 0.2 M cacodylate buffer. Then the samples were sequentially treated with 0.3% thiocarbohydrazide in 0.2 M cacodylate buffer for 10 min and 1% OsO4 in 0.2 M cacodylate buffer (pH 6,9) for 30 min. Then, samples were rinsed with 0.1 M sodium cacodylate (pH 6.9) buffer until all traces of the yellow osmium fixative have been removed, washed in de-ionized water, treated with 1% uranyl acetate in water for 1 h and washed in water again (Mironov et al., 2004; Beznoussenko et al., 2015). The samples were subsequently subjected to de-hydration in ethanol and then in acetone and embedded in Epoxy resin at RT and polymerized for at least 72 h in a 60 °C oven. Embedded samples were then sectioned with diamond knife (Diatome, Switzerland) using Leica ultramicrotome (Leica EM UC7; Leica Microsystems, Vienna). Sections were analyzed with a Tecnai 20 High Voltage EM (FEI, Thermo Fisher Scientific, Eindhoven, The Netherlands) operating at 200 kV^7^.

### Measurement of the cellular velocities and trajectories on monolayers

Coarse-grained maps of the instantaneous cellular velocities were obtained by analysing time-lapse phase-contrast movies with a custom PIV software written in MATLAB^2^. The time interval between consecutive frames was 5 min or 10 min. The interrogation window was 32X32 pixels (pixel size1.29 µm or 1.6 µm), with an overlap of 50% between adjacent windows. The number of cell comprised within one field-of-view (FOV) was typically 2500. For a given monolayer, time-lapse images from different (typically from 5 to 10) FOVs were simultaneously collected.

The instantaneous root mean square velocity *v*_*RMS*_(*t*) of a cell monolayer was computed as 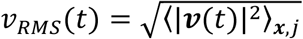, where is the instantaneous velocity vector ***v***(*t*) and ⟨ ⟩_*x,j*_ indicates an average over all grid points ***x*** (corresponding to the centers of the PIV interrogation windows) and FOVs *j*, respectively.

The instantaneous order parameter *ψ*(*t*) of a cell monolayer was computed as 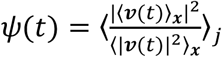. This definition is such that 0 ≤*ψ*(*t*) ≤ 1. In particular, *ψ*(*t*) = 1 only if, within each FOV, the velocity field is perfectly uniform, *i.e.* all the cells in the monolayer move with the same speed and in the same direction. On the contrary *ψ*(*t*) ≅ 0 is expected for a randomly oriented velocity field. The vectorial velocity correlation functions were calculated as 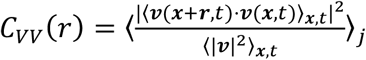. Unless otherwise stated in the main text, the temporal average ⟨ ·⟩_*t*_ was always performed over the time window comprised between 4 and 12 hours from the beginning of the image acquisition. The velocity correlation function *L*_*corr*_ is obtained by fitting *C*_*vv*_(*r*) with a stretched exponential function of the form 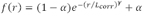. Here γ is a stretching exponent and α is an offset which is non-zero in presence of a collective migration of the monolayer.

Cellular trajectories ***r***_*m*_(*t*) were calculated by numerical integration of the instantaneous velocity field as obtained from the PIV analysis (see ref. ^8^ and reference therein). For each FOV a number of trajectories roughly corresponding to the number of cells was computed.

Mean squared displacements (MSDs) of the cells were calculated as *MSD*(Δ*t*) = ⟨ |***r***_*m*_ (*t* + Δ*t*) – ***r***_*m*_ (*t*) |^2^;⟩, where the average was performed over all the trajectories and, unless otherwise stated in the main text, in the time window comprised between 4 and 12 hours after the beginning of the experiment. In order to estimate the persistence length *L*_*pers*_ of the cellular motion the MSD curves were fitted with a function of the form *g* (*Δt*) = (*u* _0_*Δt*)^2^[1 + (*u*_0_*Δt* / *L*_*pers*_)]^-1^. This expression describes a transition between a short-time ballistic-like scaling and a long-time diffusive scaling. The transition between the two regimes takes place for *Δt ≈* 1/ *u*_0_ *L*_*pers*_, i.e. after the cell has travelled with an approximately constant velocity over a distance *≈ L*_*pers*_.

### Measurement of the cellular velocities of acini

Sequences of confocal Z stacks of 3D acini were analysed with an adapted PIV scheme in order to extract a representative value for the migration velocity, to assess the collective nature of the cellular motion and to detect the presence of a coherent rotational motion. Details about the imaging are given in the paragraph “Image acquisition”.

The geometrical centre ***x*** _*c*_ of each acinus was determined as the centroid of the corresponding 3D fluorescent intensity distribution (Z stack) *I*(*x*|*t*),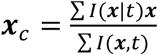, where the sum is performed over all voxels and time points. For each time point, the 3D fluorescent intensity distribution was radially projected onto the unit sphere centred in ***x*** _*c*_ leading to a sequence of 2D intensity maps *i*(*θ φ* |*t*), where *θ* and *φ* are the polar and the azimuthal angle spanning the sphere, respectively. In practice, *i*(*θ φ* |*t*) was obtained from a representation of *I*(***x*** |*t*) in spherical coordinates, after summation over the radial coordinate. For each time point, *i* is represented by a 512×128 matrix, each element covering the Cartesian product of angular intervals of constant amplitudes Δ *θ* = *π*/512 and Δ *φ*= 2*π*/128, respectively.

We performed on *i* a 2D PIV analysis as described in the previous paragraph, by treating (*θ, φ*) as Cartesian coordinates. The obtained coarse-grained velocity fields [*u* _*θ*_ (*θ,φ*|*t*), *u* _*φ*_ (*θ,φ*|*t*)] (in units of rad/hr) were then used to reconstruct the tangential velocity field ***v***(*θ, φ*) = *R*_0_ (*u* _*θ*_ (*θ, φ*|*t*)***n*** _*θ*_ +*u* _*φ*_ (*θ,φ*|*t*) sin *θ* ***n*** _*φ*_) of the acinus. Here, ***n*** _*θ*_ and ***n*** _*φ*_ are the polar and the azimuthal unit vector,respectively and 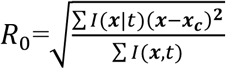 is the radius of gyration of the acinus.

The root mean squared velocity was calculated as 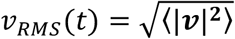, where the angular brackets indicate an average performed over the whole sphere. The presence of a pattern of global rotation was monitored by measuring the total angular momentum ***l*** = ⟨ ***r*** × ***v***⟩, where ***r*** is a unit vector spanning the whole sphere. The direction of ***l*** identifies the orientation of the axis of instantaneous rotation. The collective nature of the cellular motility is captured by the non-dimensional rotational order parameter 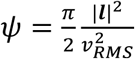. The normalization of the order parameter is such that, for a rigidly rotating sphere, *ψ* = 1, while, in the absence of coordinated motion one expects *ψ* ≅ 0.

### Kinematic and dynamical analysis of spheroids

Overall motility and internal dynamics of the spheroids were measured by analysing sequences of confocal Z stacks, according to the following procedure, implemented in a custom MATLAB^®^ code. More details about the imaging can be found in the paragraph “Image acquisition”.

We indicate with *R*(**Θ, *U***) the roto-translational operator given by the composition of a 3D rotation by an angle | **Θ**| around the axis identified by the direction of the 3D vector **Θ** and a translation of vector ***U***. *R*(**Θ, *U***)is a linear operator and its numerical implementation as a transformation between 3D matrices (Z stacks) was realized *via* the MATLAB functions *imwrap* and *affine3d*.

Let us consider two 3D stacks *I*(***x***,*t*) and *I*(***x***,*t* + Δ*t*_0_), where Δ*t*_0_ is delay between consecutive stacks. We define Ω(*t*) and ***U***(*t*) as the 3D vectors that minimize the distance *d* (namely, the variance of the difference) between *I*(***x***,*t* + Δ*t*_0_) and *R*(ω Δ*t*_0_,***u***)*I*(***x***,*t*), *d*(ω, ***u*|***t*) = ‖ *I*(***x***,*t* + Δ*t*_0_) – *R*(ω Δ*t*_0_,***u***)*I*(***x***,*t*) ‖^2^. Numerically, the minimization is performed by exploiting the MATLAB function *imregtform*. In substance, *R*(Ω(*t*) *Δt*_0_, ***U***(*t*)) is the rigid transformation that reproduces at best the changes occurred in *I*(***x***,*t*) during the time interval Δ*t* _0_. According to the definitions above, *Ω*(*t*) provides the best estimate for the instantaneous vectorial angular velocity of the spheroid, the direction of 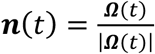 identifying the axis of instantaneous rotation. The temporal persistence of the rotational motion is captured by the orientational correlation function *C*_*n*_(Δ*t*) = ⟨ ***n***(*t*) ⟩_*t*_, where Δ*t = n*Δ*t*_0_. In order to estimate the rotational correlation time τ_*p*_, *C*_*n*_(Δ*t*) was fitted with an exponential function of the form *f*(Δ*t*) = exp(–Δ*t*/τ_*p*_).

The non-rigid part of the changes occurring within a spheroid between time *t* and *t* + Δ*t,* where Δ*t* = *n*Δ*t*_0_, is captured by the parameter: 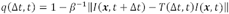, where *T*(Δ*t, t*) = *R*(*Ω*(*t* + *n*Δ*t*_0_)Δ*t*,***U***(*t* + *n*Δ*t*_0_)) *° R*(*Ω*(*t* + (*n* – 1)Δ*t*_0_) Δ*t*,***U***(*t* + (*n* – 1)Δ*t*_0_)) *°* … *°R*(*Ω*(t)Δ*t*,***U***(*t*)) is the composition of elementary roto-translations and *β* ≡ 2(⟨*I*^*^2^*^⟩ – ⟨*I*⟩^2^). The definition of *q* is such that, neglecting noise and truncation errors, *q ≅*1 if the spheroids is immobile or if it undergoes a perfectly rigid displacement and/or rotation, with no relative motion between different cells. On the contrary, one gets *q* ≅ 0 when almost all the cells have performed positional rearrangements on a length scale comparable with their size, leading to a substantial change in the local structure^9^. We consider in particular the so-called overlap parameter *Q*, obtained as a temporal average of *q* :*Q*(Δ*t*) = ⟨*q* (Δ*t,t*)_t._ By fitting the decay of *Q* with an exponential function 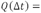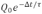 we can extract an estimate of the characteristic correlation time τ after which an almost complete change in the cellular configuration has occurred.

### PIV analysis on ex vivo tumour slices

Maps of the instantaneous cellular velocities were obtained by analysing time-lapse movies by performing a PIV analysis using the MATLAB (Release R2017b The MathWorks, Inc., Natick, Massachusetts, United States) MPIV toolbox (http://www.oceanwave.jp/softwares/mpiv/)^10^ with the correlation algorithm and an interrogation window of 24 pixels X 24 pixels (1 pixel = 1.4 um).

The analysis was performed on 3 independent experiments per condition on border sections of the tumour (for a total of 5 field of view per condition).

The instantaneous root mean square velocity v_RMS_(t) of a single Field of View was calculated as:

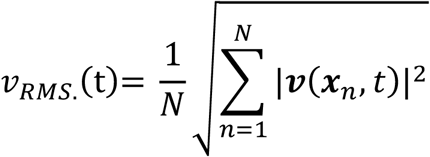

where N is the number of grid points in the field of view and ***v***(***x****_n_, t*) is the instantaneous velocity at the *n*th grid point ***x***,_*n*_.

### Statistical analysis

Student’s unpaired and paired t-test was used for determining the statistical significance. Significance was defined as **\*** P < 0.05;**P < 0.001 and ****P < 0.0001.Statistic calculations were performed with GraphPad Prism Software. Data are expressed as mean±SD, unless otherwise indicated.

